# Insights into the involvement of spliceosomal mutations in myelodysplastic disorders from an analysis of SACY-1/DDX41 in *Caenorhabditis elegans*

**DOI:** 10.1101/2019.12.23.886804

**Authors:** Tatsuya Tsukamoto, Micah D. Gearhart, Seongseop Kim, Gemechu Mekonnen, Caroline A. Spike, David Greenstein

## Abstract

Mutations affecting spliceosomal proteins are frequently found in hematological malignancies, including myelodysplastic syndromes and acute myeloid leukemia. DDX41/Abstrakt is a metazoan-specific spliceosomal DEAD-box RNA helicase found to be recurrently mutated in inherited myelodysplastic syndromes and in relapsing cases of acute myeloid leukemia. The genetic properties and genomic impacts of disease-causing missense mutations in DDX41 and other spliceosomal proteins have been uncertain. Here we conduct a comprehensive molecular genetic analysis of the *C. elegans* DDX41 ortholog, SACY-1. Our results reveal general essential functions for SACY-1 in both the germline and the soma, as well as specific functions affecting germline sex determination and cell cycle control. Certain *sacy-1/DDX41* mutations, including the R525H human oncogenic variant, confer antimorphic activity, suggesting that they compromise the function of the spliceosome. Consistent with these findings, *sacy-1* exhibits synthetic lethal interactions with several spliceosomal components, and biochemical analyses suggest that SACY-1 is a component of the *C. elegans* spliceosome. We used the auxin-inducible degradation system to analyze the impact of SACY-1 on the transcriptome using RNA sequencing. SACY-1 depletion impacts the transcriptome through splicing-independent and splicing-dependent mechanisms. The observed transcriptome changes suggest that disruption of spliceosomal function induces a stress response. Altered 3’ splice site usage represents the predominant splicing defect observed upon SACY-1 depletion, consistent with a role for SACY-1 as a second-step splicing factor. Missplicing events appear more prevalent in the soma than the germline, suggesting that surveillance mechanisms protect the germline from aberrant splicing.

**Author Summary:** Mutations affecting spliceosomal proteins are frequently found in hematological malignancies. DDX41/Abstrakt is a metazoan-specific spliceosomal DEAD-box RNA helicase recurrently mutated in inherited and relapsing myelodysplastic syndromes and acute myeloid leukemia. The genetic properties and genomic impacts of disease-causing mutations in spliceosomal proteins have been uncertain. Here we conduct a comprehensive molecular genetic analysis of the *C. elegans* DDX41 ortholog, SACY-1. Our results reveal that multiple *sacy-1/DDX41* missense mutations, including the R525H human oncogenic variant, exhibit antimorphic activity that likely compromises the function of the spliceosome. The genomic consequences of SACY-1 depletion include splicing-splicing-independent and splicing-dependent alterations in the transcriptome.

## INTRODUCTION

Mutations affecting components of the spliceosome are frequently found in hematological malignancies, including myelodysplastic syndromes (MDS; Yoshida *et al*. 2011; reviewed by Yoshida and Ogawa 2014; Coltri *et al*. 2019), which comprise a heterogeneous set of myeloid neoplasms characterized by anemia and cytopenia that progress to acute myeloid leukemia (AML) to varying degrees (Tefferi and Vardiman 2009). The genetic properties and genomic impacts of disease-causing missense mutations in DDX41 and other spliceosomal proteins have been uncertain. Nonetheless, mutations affecting spliceosomal components are predictive of poor clinical outcomes in AML patients (Papaemmanuil *et al*. 2016). Exactly how mutations in spliceosomal components contribute to malignancy is uncertain, but an attractive model is that aberrant splicing may interrupt tumor suppressor activity. Importantly, genome sequencing data in patients is currently being used in the clinic to generate personalized prognoses, with the idea of optimally targeting existing therapies and generating new treatment strategies (Grinfeld *et al*. 2018). One potential therapeutic approach under development is the discovery of splicing inhibitors (Effenberger *et al*. 2017; Kim and Abdel-Wahab 2017; DeNicola and Tang 2019). Although mutations affecting several spliceosomal proteins appear to be beneficial to tumor cells, excessive splicing abnormalities are likely to be lethal to all cells. Splicing inhibitors have been demonstrated to target tumor cells with splicing mutations by inducing excessive splicing abnormalities, but cells with intact splicing machinery appear to be resistant to these agents (Seiler *et al*. 2018). In fact, several new splicing inhibitors are currently in clinical trials.

The spliceosomal components frequently affected in MDS include the biochemically well-defined factors SF3B1, SRSF2, and U2AF1 (Yoshida *et al*. 2011; reviewed by Yoshida and Ogawa 2014). More recent studies have implicated DDX41 (Ding *et al*. 2012; Lewinsohn *et al*. 2015; Polprasert *et al*. 2015; Cardoso *et al*. 2016; Li *et al*. 2016; Diness *et al*. 2018; reviewed by Maciejewski *et al*. 2017), a DEAD-box RNA helicase highly conserved in metazoans, whose precise biochemical function in the spliceosome is less well understood. DDX41 appears to be specifically recruited to the catalytically active C complex (Jurica *et al*. 2002; Bessonov *et al*. 2008), which performs the second step of splicing in which the 5’ and 3’ exons are ligated and an intronic lariat is released. DDX41 is one of many spliceosomal proteins specific to metazoans and not found in budding yeast (Bessonov *et al*. 2008).

Whole genome sequencing studies suggest that *DDX41* mutations are associated with hematological malignancies that are considered to be different clinical entities. For example, examination of clonal evolution of relapsed AML cases identified *DDX41* as one of several genes found to be mutated in secondary tumors but not the primary tumors, suggesting it plays a role as a tumor suppressor and might be causative for disease progression (Ding *et al*. 2012). By contrast, studies of familial acute myeloid leukemia syndromes suggest that preexisting germline *DDX41* mutations in trans to newly arising somatic mutations cause the development of hematological malignancies (Polprasert *et al*. 2015; Cardoso *et al*. 2016; Lewinsohn *et al*. 2016; Li *et al*. 2016). Germline biallelic *DDX41* missense mutations were recently reported in two siblings that exhibited intellectual disability, psychomotor delays, and facial and skeletal dysmorphologies, with one sibling presenting with childhood leukemia (Diness *et al*. 2018). Other work suggests that DDX41 might be a multifunctional protein; in addition to its nuclear function in RNA splicing, it has been suggested to function as a cytoplasmic DNA sensor in a signaling pathway in the cytoplasm that detects infecting double stranded DNA and initiates an antiviral interferon response (Zhang *et al*. 2011; Parvatiyar *et al*. 2012; Stavrou *et al*. 2015, 2018; reviewed by Jiang *et al*. 2017). However, more recent work suggests that cyclic GMP-AMP synthase (cGAS) functions as the major DNA sensor and is several orders of magnitude more effective in inducing interferon beta synthesis than DDX41 (Sun *et al*. 2013). Two studies, one of DDX41 and another of its *Drosophila* ortholog, Abstrakt, suggested a role in regulating translation of the cyclin-dependent kinase inhibitor p21^WAF1/CIP1^ (Peters *et al*. 2017) and the Inscuteable protein (Irion *et al*. 2004), respectively, though the exact mechanism for these activities has not been elucidated and indirect effects acting at the level of splicing were not addressed in these studies.

To better understand the highly conserved functions of *DDX41*, we undertook a comprehensive molecular genetic analysis of its ortholog, *sacy-1* in the nematode *Caenorhabditis elegans*. Our prior studies identified the DEAD-box helicase SACY-1 as a negative regulator of oocyte meiotic maturation functioning in the germline upstream of the TIS11 CCCH zinc-finger RNA-binding proteins OMA-1 and OMA-2 (Kim *et al*. 2012). Genetic analysis also established roles for SACY-1 in regulating the hermaphrodite sperm-to-oocyte switch and in preventing necrotic cell death of gametes. Genetic experiments further suggested an essential role for *sacy-1(+)* in early embryos and larvae that appeared to be maternally rescued. At the time of our original study, searchable databases of the scientific literature had not yet annotated DDX41 (or its *Drosophila* ortholog, Abstrakt) as spliceosomal components identified by proteomics. We therefore did not recognize that SACY-1 was likely involved in splicing.

In this study, we undertook a comprehensive molecular genetic analysis of SACY-1’s functions in *C. elegans*. Our results are most consistent with an essential role for SACY-1 in spliceosome function. Further, our genetic results reveal that certain *sacy-1* mutations appear to confer a dosage-sensitive antimorphic activity, most consistent with the possibility that they compromise the function of the spliceosome by perturbing the function of other spliceosomal proteins. The oncogenic R525H mutation in human DDX41 was introduced into the *C. elegans* genome using CRISPR-Cas9 genome editing and found to exhibit weak antagonistic activity. Consistent with these findings, *sacy-1* exhibits genetic interactions with associated spliceosomal components, and biochemical analyses suggest that SACY-1 is a component of the *C. elegans* spliceosome. Depletion of SACY-1 in the germline or soma was found to have major impacts on the transcriptome through splicing-independent and splicing-dependent mechanisms. Alterations in 3’ splice site selection represent the most prevalent changes in splicing patterns observed following SACY-1 depletion, consistent with its function as a component of the spliceosomal C complex. Missplicing events are more prevalent upon SACY-1 depletion in the soma than in the germline, leading us to suggest that surveillance mechanisms protect the germline from aberrant splicing. The observed gene expression changes observed after SACY-1 depletion suggest that perturbations of spliceosomal function might induce a stress response. Our results, taken together with a recent study of *sftb-1/SF3B1* (Serrat *et al*. 2019), highlight the potential of the *C. elegans* system for examining the molecular genetic properties of disease-causing mutations affecting highly conserved components of the spliceosome.

## MATERIALS AND METHODS

### C. elegans strains and genetic analysis

The genotypes of strains used in this study are reported in Supporting Information, Table S1. Genes and mutations are described in WormBase (www.wormbase.org<u>; Harris *et al*. 2013</u>) or in the indicated references. Culture and genetic manipulations were conducted at 20°C unless specified otherwise. The following mutations were used: LGI– *fog-1(q253*ts), *dpy-5(e61)*, *gld-1(tn1478)*, *unc-13(e51)*, *unc-13(e1091)*, *lin-41(n2914)*, *lin-41(tn1541[gfp::tev::s-tag::lin-41])*, *sacy-1(tm5503)*, *sacy-1(tn1385)*, *sacy-1(tn1479)*, *sacy-1(tn1480)*, *sacy-1(tn1481*Mog), *sacy-1(tn1482)*, *sacy-1(tn1602)*, *sacy-1(tn1603)*, *sacy-1(tn1604)*, *sacy-1(tn1605)*, *sacy-1(tn1606)*, *sacy-1(tn1607)*, *sacy-1(tn1608)*, *sacy-1(tn1609)*, *sacy-1(tn1610)*, *sacy-1(tn1611)*, *sacy-1(tn1612)*, *sacy-1(tn1615)*, *sacy-1(tn1616)*, *sacy-1(tn1617)*, *sacy-1(tn1632[3xFLAG::PreScission protease site::gfp::tev::s-tag::sacy-1])*, *sacy-1(tn1880[aid::gfp::tev::myc::sacy-1])*, and *sacy-1(tn1887)*; LGII–*tra-2(e2020)*, *ieSi57[eft-3p::TIR1::mRuby::unc-54 3’UTR + Cb unc-119(+)], ieSi64[gld-1p::TIR1::mRuby::gld-1 3’UTR + Cb unc-119(+)]*; LGIII–*unc-119(ed3)*; LGIV–*unc-24(e138)*, *fem-3(e1996)*, and *dpy-20(e1282)*; LGV–*acy-4(ok1806)*, *her-1(hv1y101)*, *emb-4(sa44)*, *unc-51(e369)*, and *fog-2(oz40)*. The following rearrangements were used: *hT2[bli-4(e937) let-?(q782) qIs48]* (I;III), *tmC18[dpy-5(tmIs1236) + pmyo-2::mCherry]* I (Dejima *et al*. 2018), *mIn1[dpy-10(e128) mIs14]* II, and *tmC12[egl-9(tmIs1194) + pmyo-2::Venus]* V (Dejima *et al*. 2018). The following transgenes were used: *tnEx37[acy-4(+)* + *sur-5::gfp]*, *tnEx159[gfp:sacy-1 + pDPMM0016B(unc-119(+))]*.

**Table 1.** RNAi of *sacy-1* enhancer loci increase the penetrance of germline or lethal phenotypes in *sacy-1(tn1385)* reduction-of-function mutants

| RNAi<sup>a</sup> | Genotype | Sterile (%)<sup>b</sup> | Gamete<br>degeneration<sup>b</sup><br>(%) | Embryonic lethal (%)<sup>c</sup> |
| --- | --- | --- | --- | --- |
| L4440 (control) | Wild type | 0<br>(n=338) | 0<br>(n=338) | 1<br>(n=674) |
|  | *sacy-1(tn1385)* | 0<br>(n=256) | 0<br>(n=256) | 1<br>(n=502) |
| *mog-2* (II-3D16) | Wild type | 4<br>(n=172) | 0<br>(n=54) | 7<br>(n=574) |
|  | *sacy-1(tn1385)* | 43<br>(n=254) | 47<br>(n=72) | 95<br>(n=426) |
| *Y111B2A.25*<br>(III-6G22) | Wild type | 22<br>(n=230) | 1<br>(n=94) | 84<br>(n=463) |
|  | *sacy-1(tn1385)* | 95<br>(n=272) | 16<br>(n=110) | 93<br>(n=42) |
| *cacn-1*<sup>d</sup> (II-9E09) | Wild type | 100<br>(n=164) | 35<br>(n=40) | ND |
|  | *sacy-1(tn1385)* | 100<br>(n=224) | 81<br>(n=52) | ND |
| *emb-4* (V-12E12) | Wild type | 2<br>(n=184) | 0<br>(n=110) | 12<br>(n=337) |
|  | *sacy-1(tn1385)* | 89<br>(n=176) | 42<br>(n=82) | N.D. |
| *emb-4* (V-12E14) | Wild type | 4<br>(n=288) | 0<br>(n=96) | 13<br>(n=421) |
|  | *sacy-1(tn1385)* | 91<br>(n=202) | 80<br>(n=120) | ND |
| *emb-4* (V-12E16) | Wild type | 4<br>(n=210) | 0<br>(n=96) | 9<br>(n=433) |
|  | *sacy-1(tn1385)* | 91<br>(n=326) | 87<br>(n=140) | ND |

<sup>a</sup>RNAi clones showing genetic interactions with *sacy-1* are listed with the target gene name in italics and the location of the clone in the RNAi library in parentheses. The identity of clones was verified by DNA sequencing.
<sup>b</sup>Sterility and gamete degeneration were scored by DIC microscopy on the first day of adulthood 24 hours post-L4 at 22°C. Gonad arms were scored as sterile if they did not produce embryos and exhibited defects in gametogenesis. Number of gonad arms scored is reported.
<sup>c</sup>Embryonic lethality was measured by conducting daily egg lays over the reproductive lifespan and determining the number of embryos that failed to hatch by 48 hours after egg laying. The number of embryos scored is reported. <sup>d</sup>*cacn-1* was not initially identified as an enhancer of *sacy-1* during the genome-wide RNAi screen because *cacn-1(RNAi)* results in complete sterility in both *sacy-1(tn1385)* and wild-type animals. However, we determined that RNAi to the *Y111B2A.25* pseudogene likely targets *cacn-1* (see text for details). ND, not determined.

For the analysis of genetic interactions between *sacy-1(tn1481)* and *fem-3(e1996)*, non-Unc non-Dpy non-GFP animals from *sacy-1(tn1481)/hT2[bli-4(e937) let-?(q782) qIs48]; fem-3(e1996)/unc-24(e138) dpy-20(e1282)* were individually cultured and scored for germline phenotypes. Following scoring, the *fem-3* genotype of each animal was scored by conducting PCR with primers *fem-3* F2 and *fem-3* R2 and sequencing the products.

To map the cold-sensitive (15°C) and temperature-sensitive (25°C) phenotypes of *sacy-1(tn1480)*, 34 Unc non-Dpy recombinants were obtained from *sacy-1(tn1480)/dpy-5(e61) unc-13(e1091)* heterozygotes. The recombinant chromosomes were bred to homozygosity and scored for the presence or absence of the *sacy-1(tn1480)* mutation by conducting PCR with primers H27M09.1F1 and H27M09.1R4 and sequencing purified PCR products with primer H27M09.1F2. We found that 7 of the 34 recombinants contained *sacy-1(tn1480)* and were cold-sensitive and temperature-sensitive. By contrast, 27 recombinants were *sacy-1(+)* and grew at 15°C and 25°C. These data indicate that *sacy-1(tn1480)* mutation is inseparable from the cold-sensitive and temperature-sensitive phenotypes (e.g., within ∼0.06 map units). In addition, 32 Dpy non-Unc recombinants were selected. Interestingly, all the homozygous recombinants were fertile at both 15°C and 25°C, including the 22 recombinants that contained the *sacy-1(tn1480)* mutation. Although these *dpy-5(e61) sacy-1(tn1480)* recombinants grew at 15°C and 25°C, they produced appreciable numbers of dead embryos and grew more slowly than their *sacy-1(+)* counterparts. This result suggests that the *dpy-5(e61)* mutation suppresses the cold-sensitive and temperature-sensitive phenotypes of *sacy-1(tn1480)*. Previous work has shown that mutant alleles of collagen genes can suppress temperature-sensitive mutations in other gene products possibly by triggering a stress response (Levy *et al*. 1993; Maine and Kimble 1993; Nishiwaki and Miwa 1998). That *dpy-5(e61)* suppresses *sacy-1(tn1480)* was further shown by constructing *dpy-5(e61) sacy-1(tn1480)/sacy-1(tn1480) unc-13(e1091)* heterozygotes (n=30), of which 20 exhibited the *sacy-1(tn1480)* sperm-defective phenotype at 25°C and 10 were fertile. Thus, *dpy-5(e61)* exhibits semidominance for its effects on body morphology and for suppression of *sacy-1(tn1480)*. To examine the dominant Him phenotype of *sacy-1(tn1480)* and its interaction with *sacy-1(tn1887)*, we compared the percentage of males produced at 25°C by *dpy-5(e61)/sacy-1(tn1480) unc-13(e1091)* and *dpy-5(e61) sacy-1(tn1887)/sacy-1(tn1480) unc-13(e1091)* heterozygotes.

### RNA interference

Genome-wide RNA interference (RNAi) screening employed the Ahringer feeding library (Kamath *et al*. 2003) using the RNAi culture media described by Govindan *et al*. (2006) at 22°C. The empty vector L4440 was used as a control. The identity of RNAi clones was verified by DNA sequencing. Gene-specific RNAi was performed by placing gravid hermaphrodites on RNAi medium seeded with double-stranded RNA (dsRNA)-expressing *E. coli* (Timmons and Fire 1998). The gravid hermaphrodites were immediately treated with 20% bleach to release the F1 embryos. Phenotypes were assessed 3–4 days later. For quantification of phenotypes, sterility and gamete degeneration were scored in the F1 generation, and embryonic lethality was scored in the F2 generation produced by the RNAi-treated F1 animals.

### Immunofluorescence, fluorescent labeling, and microscopy

Dissected gonads were fixed in 3% paraformaldehyde as described (Rose *et al*. 1997). Fixed gonads were stained with rabbit anti-RME-2 antibody (Grant and Hirsh 1999; kindly provided by B. Grant, Rutgers University, 1:50), a mixture of two purified mouse monoclonal anti-MSP antibodies (Kosinski *et al*. 2005, each at 1:300), rabbit anti-phospho-histone H3 (Ser10) antibody (Millipore, 1:400). Secondary antibodies were Alexa 488-conjugated donkey anti-rabbit antibodies (Jackson ImmunoResearch, 1:500) and Cy3-conjugated goat anti-mouse antibodies (Jackson ImmunoResearch, 1:500). 4’, 6-diamidino-2-phenylindole (DAPI) was used to detect DNA. DIC and fluorescent images were acquired on a Zeiss motorized Axioplan 2 microscope with either a 40x Plan-Neofluar (numerical aperture 1.3) or a 63x Plan-Apochromat (numerical aperture 1.4) objective lens using a AxioCam MRm camera and AxioVision software (Zeiss).

### Genome editing

CRISPR-Cas9 genome editing used pRB1017 to express single guide RNA (sgRNA) under control of the *C. elegans* U6 promoter (Arribere *et al*. 2014). The sequences of all oligonucleotides used are listed in Table S2. To generate sgRNA clones, annealed oligonucleotides were ligated to *Bsa*I-digested pRB1017 plasmid vector, and the resulting plasmids were verified by Sanger sequencing. pDD162 served as the source of Cas9 expressed under control of the *eef-1A.1/eft-*3 promoter (Dickinson *et al*. 2013). Indels were targeted to exon 2 of *sacy-1* using *sacy-1* sgRNA7 (pCS520). The injection mix contained pCS520 (25 ng/μl), pDD162 (50 ng/μl), and *Pmyo-2::tdTomato* (4 ng/μl). *sacy-1(tn1602– tn1612)* were recovered from injections into DG3913 *lin-41(tn1541[gfp::tev::s-tag::lin-41])* and *sacy-1(tn1615–1617)* were recovered from injections into the wild type (strain N2).

**Table 2.** Genetic interactions between *sacy-1* and *emb-4*

**A. Enhancement of germline and somatic *sacy-1* mutant defects**
| Genotype | Vulval<br>rupture<br>(%) | Sterile<br>and Pvl<sup>a</sup><br>(%) | Sterile<sup>a</sup><br>(%) | Fertile<br>(%) |
| --- | --- | --- | --- | --- |
| *sacy-1(tm5503)* (n=284) | 3 | 1 | 96 | 0 |
| *sacy-1(tm5503)/+; emb-4(sa44)<sup>b</sup>* (n=205) | 0 | 0 | 0 | 100 |
| *sacy-1(tm5503); emb-4(sa44)<sup>b</sup>* (n=242) | 83 | 16 | 1 | 0 |
| *sacy-1(tn1385)* (n=278) | 0 | 0 | 0 | 100 |
| *sacy-1(tn1385)/+; emb-4(sa44)<sup>b</sup>* (n=201) | 0 | 0 | 0 | 100 |
| *sacy-1(tn1385); emb-4(sa44)<sup>b,c</sup>* (n=144) | 36 | 4 | 3 | 57<sup>d</sup> |

<sup>a</sup>Sterile animals exhibit the *sacy-1(lf)* gamete degeneration phenotype.
<sup>b</sup>The *hT2(qIs48)* balancer chromosome, which is dominantly marked with GFP, was used to differentiate between *sacy-1(tm5503)* heterozygotes and homozygotes.
<sup>c</sup>The progeny of *sacy-1(tn1385)/hT2(qIs48); emb-4(sa44)* hermaphrodites; the balancer chromosome provides maternal *sacy-1(+)* function. The fertile F1 progeny of these animals are maternal-effect lethal, see Table 2B, below.
<sup>d</sup>These adult hermaphrodites produce a majority of embryos that fail to hatch, see Table 2B.

**B. Enhancement of embryonic lethality**
| Genotype<sup>a</sup> | Embryonic lethal (%) | L1 lethal (%) | Viable (%) |
| --- | --- | --- | --- |
| *sacy-1(tn1385)* (n=949) | 1 | 0 | 99 |
| *emb-4(sa44)* (n=1348) | 4 | 0 | 96 |
| *sacy-1(tn1385); emb-4(sa44)<sup>b</sup>* (n=620) | 97 | 3 | 0 |

<sup>a</sup>The number of embryos examined.
<sup>b</sup>The F1 progeny of fertile *sacy-1(tn1385); emb-4(sa44)* parents derived from the *sacy-* *1(tn1385)/+; emb-4(sa44)* heterozygotes analyzed in Table 2A above.

An N-terminal *gfp* fusion to endogenous *sacy-1*, *sacy-1(tn1632[3xflag::PreScission protease site::gfp::tev::s-tag::sacy-1]),* was constructed using *sacy-1* sgRNA1 (pCS486) and a repair template generated by conducting the PCR with oligonucleotide primers *sacy-1* 5HAF and *sacy-1* 3HAR, using a *gfp::sacy-1::tev::s-tag* recombineered fosmid (SK212; Kim *et al*. 2012) as template. Genome editing employed the *dpy-10* co-conversion method (Arribere *et al*. 2014). The injection mix contained pJA58 (7.5 ng/μl), AF-ZF-827 (500 nM), pCS486 (50 ng/μl), repair template (50 ng/μl), and pDD162 (50 ng/μl) and was injected into wild-type worms. Correct targeting was verified by conducting PCR with primer pairs GFP_7215 and H27M09.1_R5 and GFP_1094R and H27M09.1_seqF1 followed by DNA sequencing.

An N-terminal auxin-inducible degron (aid) fusion to *sacy-1*, *sacy-1(tn1880[aid::gfp::myc::sacy-1]),* was constructed using *sacy-1* sgRNA1 and a repair template generated by conducting the PCR with oligonucleotide primers *sacy-1* AID5F and *sacy-1* AID3R using a *wee-1.3::aid::gfp::myc* clone (pCS575, C. Spike, unpublished results) as template. The injection mix was prepared as described above and was injected into CA1352 worms. *sacy-1* (*sacy-1(tn1880[aid::gfp::myc::sacy-1])* was identified by screening the progeny of 414 F1 Roller animals for GFP fluorescence. Correct targeting was verified by conducting PCR with primer pairs GFP_R1 and H27M09.1_F5 and GFP_F1 and H27M09.1_R5 followed by DNA sequencing.

The R525H mutation in DDX41 was imported into *C. elegans* (e.g., SACY-1[R534H]) using genome editing (Paix *et al*. 2014) with *sacy-1* sgRNA11 and *sacy-1* sgRNA12 and a single-stranded repair oligonucleotide (*sacy-1* GM1), which introduces the R534H mutation and two synonymous changes to alter the protospacer adjacent motif and to facilitate screening using an introduced *Ava*I restriction site. The injection mix contained pJA58 (7.5 ng/μl), AF-ZF-827 (500 nM), *sacy-1* sgRNA11 (25 ng/μl), *sacy-1* sgRNA12 (25 ng/μl), *sacy-1* GM1 (500 nM), and pDD162 (50 ng/μl) and was injected into wild-type worms. Edited loci were verified by PCR and DNA sequencing using primers sacy-1 seq F1 and sacy-1 seq R1.

### Antibody production, purification, and western blotting

*sacy-1* cDNA sequences were cloned into the *E. coli* expression vector pMal-c2 to create an inducible fusion protein wherein maltose binding protein was fused to amino acids 411–578 of SACY-1 (MBP::SACY-1(411–578)). MBP::SACY-1(411–578) was column and gel-purified and used to immunize rabbits. Immunizations and sera collection were performed using standard protocols (Cocalico Biologicals, Inc., Reamstown, PA). Rabbit antibody (R217) was affinity purified and was suitable in western blots with partially purified SACY-1 preparations. Hybridoma cell lines producing anti-GFP monoclonal antibodies 12A6 and 4C9 (Sanchez *et al*. 2014) were obtained from the Developmental Studies Hybridoma Bank and prepared as described (Tsukamoto *et al*. 2017). Proteins were separated using NuPAGE 4-12% Bis-Tris gels (Invitrogen) and visualized after western blotting. Blots were blocked with 5% nonfat dried milk. Primary antibodies used to detect proteins were affinity-purified rabbit anti-SACY-1(411–578) R217 antibody (100 ng/ml) and rabbit anti-GFP NB600-308 antibody (Novus Biologicals; 250 ng/ml). The secondary antibody used for western blots was peroxidase-conjugated donkey anti-rabbit antibody (Jackson ImmunoResearch; 1:30,000). Detection was performed using SuperSignal West Femto Maximum Sensitivity Substrate (Thermo Scientific).

### SACY-1 tandem affinity purification

Tandem affinity purification of SACY-1 was conducted using strains DG4068 and DG4070 using modifications of a previously described protocol (Tsukamoto *et al*. 2017). Immunopurified proteins were precipitated with 16.7% trichloroacetic acid (TCA), washed with acetone at –20°C, and briefly separated on a 12% NuPAGE Bis-Tris gel, stained with Colloidal Blue Staining Kit (Invitrogen). Lanes were subdivided into eight gel slices and mass spectrometry was performed at the Taplin Biological Mass Spectrometry Facility (Harvard Medical School) using an LTQ Orbitrap Velos Pro ion-trap mass spectrometer (Thermo Fisher Scientific). Protein identification used the Sequest software program (Thermo Fisher Scientific) to match the fragmentation pattern of tryptic peptides to the *C. elegans* proteome. The data were filtered to a 1–2% peptide false discovery rate. File S1 reports the mass spectrometry results and the additional filtering criteria for identifying non-specific interactions.

### RNA sequencing

The auxin-inducible degradation system (Zhang *et al*. 2015) was used to deplete SACY-1 using strain backgrounds in which TIR1 was expressed in the germline (CA1352) or soma (CA1200). Experimental (DG4700 and DG4703) and control strains (CA1352 and CA1200) were grown on peptone-enriched nematode growth medium with NA22 as a food source. Embryos were isolated by alkaline hypochlorite treatment (20% bleach and 0.5 N NaOH), washed in M9 buffer and allowed to hatch overnight in the absence of food. For each of three biological replicates, 60,000 L1-stage larvae were cultured on two 150-by 15-mm petri dishes containing peptone-enriched medium with OP50. The worms were grown to the young adult stage and harvested by washing off the plates with M9 and then placed on fresh plates containing peptone-enriched medium and 2 mM auxin seeded with OP50. Plates were cultured in the dark at 20°C for 24 hours. The worms were then harvested and washed with M9 repeatedly to reduce the presence of *E. coli*. Total RNA was isolated using TRIzol LS Reagent (Invitrogen, Carlsbad, CA) and the RNAeasy Micro Kit (QIAGEN, Valencia, CA). Poly(A)+ RNA was selected from 1 μg of total RNA using the NEBNext Ultra Kit (New England Biolabs, Ipswitch, MA). Libraries were prepared and sequenced by Genewiz (South Plainfield, NJ). Paired-end reads of 150 base pairs (bp) were obtained on an Illumina HiSeq4000 instrument with an average depth greater than 31 million reads per sample.

### Bioinformatics

After trimming adapters with Trim Galore (v0.6.0) and cutadapt (v1.18), reads were assessed for quality with FastQC (v0.11.8), mapped to the WBcel235/ce11 genome with STAR (v2.7.2a) guided by gene annotations defined in Ensembl (release 97) and sorted and indexed with samtools (v1.7). Gene-level abundance was estimated for Ensembl defined annotations using the featureCounts function in the Bioconductor package Rsubread (v1.28.1). An average of 28 million high-quality (MAPQ > 55) reads mapped to annotated genes within each sample. Principal component analysis and inspection of 5’ vs 3’ read coverage indicated that one soma control sample (CA1200-2) contained degraded RNA and was excluded from further analysis. Differential gene expression of Ensembl defined genes was determined using DESeq2 (v1.26.0). P values were adjusted for multiple test correction using Benjamini–Hochberg procedure. The fold change, adjusted p values, the mean number of counts across samples and the number of complementary DNA fragments per kilobase of transcript per million mapped reads (FPKM) were used to define differentially expressed genes. Gene ontology (GO) data were obtained from WormBase release WS273 and analyzed taking length bias into account using the Goseq (v1.38.0) package. Novel transcripts in each of the high quality samples and in the previously published GSE57109 (Ortiz *et al*. 2014) dataset were identified using StringTie (v2.0.4) and merged together with the Ensembl annotations to generate a comprehensive annotation set. These annotations were used with RMATS (v4.0.2 turbo) to determine statistically significant differences for splicing events between conditions [expressed as false discovery rates (FDRs)]. Coverage data was visualized with Gviz (v 1.30.0). A custom R script with details for the analysis and figure generation is available at https://github.com/micahgearhart/sacy1.

### Data availability

Strains and reagents are available upon request. RNA sequencing data have been deposited in NCBI’s Gene Expression Omnibus and are accessible through accession number XXXX.

## RESULTS

### Synthetic lethal interactions between a sacy-1 reduction-of-function allele and several genes encoding spliceosomal proteins

In prior studies, we recovered reduction-of-function (rf) *sacy-1* mutant alleles as <u>s</u>uppressors of *<u>acy</u>-4* sterility (Kim *et al*. 2012). ACY-4 is an adenylate cyclase that functions in the gonadal sheath cells to promote oocyte growth and meiotic maturation (Govindan *et al*. 2009; Nadarajan *et al*. 2009). *acy-4* null mutants are sterile, whereas *sacy-1(*rf*); acy-4(0)* mutants are fertile, and this suppression was shown to involve reduced *sacy-1* function in the germline (Kim *et al*. 2012). *sacy-1(*rf) mutants were also found to suppress the self-sterility of *fog-2* null mutants that results from a failure to produce sperm. A sperm-to-oocyte switch governs hermaphrodite germline sex determination. In wild-type hermaphrodites, the first differentiating germ cells produce sperm in the L4 larval stage, whereas, later differentiating germ cells exclusively produce oocytes in the adult stage. The suppression of *fog-2* sterility suggested that *sacy-1* has a function to promote the oocyte fate. Analysis of the *sacy-1(tm5503)* deletion allele, which is a likely null allele (see below), indicated that *sacy-1* also functions to prevent necrotic degeneration of sperm and oocytes and is required for fertility of hermaphrodites and males (Kim *et al*. 2012).

To better understand the function of *sacy-1* in multiple germline processes, we conducted a genome-wide RNAi screen for loci that enhance *sacy-1* mutant phenotypes, such as sterility or embryonic lethality. Specifically, we screened for loci which caused severe phenotypes when knocked down by RNAi in the *sacy-1(tn1385*rf) genetic background but not in the wild type. We screened 18,101 RNAi clones from the Ahringer RNAi library and identified five clones that enhanced *sacy-1* mutant phenotypes when they are knocked down (Table 1). The five RNAi clones target the transcripts of three genes (Table 1): *mog-2* (one clone), *Y111B2A.25* (one clone), and *emb-4* (three clones). To test whether *sacy-1* expression and/or localization is affected by RNAi of *mog-2*, *Y111B2A.25*, or *emb-4*, we conducted RNAi of these genes in *sacy-1(tm5503)* mutant animals expressing the rescuing *gfp::sacy-1* transgene (*tnEx159*). In no case did we observe that an RNAi treatment altered the expression or localization of the GFP::SACY-1 transgene; the expression level and predominant nuclear localization of GFP::SACY-1 after RNAi was similar to that of the control animals (S. Kim and D. Greenstein, unpublished results). This result suggests that the RNAi treatments enhance *sacy-1(tn1385)* mutant phenotypes through effects independent of SACY-1 expression and localization.

*mog-2(RNAi)* induces a higher penetrance of sterility, gamete degeneration, and embryonic lethality in the *sacy-1(tn1385)* mutant genetic background in comparison to the wild type (Table 1; Figure S1). *mog-2* encodes the U2 snRNP protein A’ (Zanetti *et al*. 2011), which is a constitutive component of the spliceosome (Jurica *et al*. 2002; Bessonov *et al*. 2008, 2010; Herold *et al*. 2009).

Similarly, *Y111B2A.25(RNAi)* specifically enhances the penetrance of multiple *sacy-1* mutant phenotypes, including sterility, gamete degeneration, and embryonic lethality (Table 1; Figure S1). *Y111B2A.25* is annotated as a pseudogene (Agarwal *et al*. 2010; www.wormbase.org). *Y111B2A.25* is part of an operon, and the expressed sequence tag (EST) data show that the *Y111B2A.25* locus is transcribed, but the transcript lacks protein-coding ability. In *C. elegans* ∼40 bp of sequence identity is sufficient to induce off-target RNAi effects (Rual *et al*. 2007). Use of the Basic Local Alignment Search Tool (BLAST) indicates that *Y111B2A.25(RNAi)* might target the *cacn-1* locus, which encodes a spliceosomal protein and shares ∼200 bp of sequence identity with *Y111B2A.25*. To test whether the enhanced sterility induced by *Y111B2A.25(RNAi)* in the *sacy-1(tn1385)* genetic background might be explained by an off-target effect to *cacn-1*, we conducted *cacn-1(RNAi)* and found that the *cacn-1(RNAi)* induces complete sterility in both the *sacy-1(tn1385)* and wild-type animals (Table 1 and Figure S1). Interestingly, under the *cacn-1(RNAi)* condition, the *sacy-1(tn1385)* animals show additional phenotypes, such as high penetrance of a protruding vulva (Pvl) phenotype and slow growth compared to the wild type, suggesting a genetic interaction between *cacn-1* and *sacy-1* that might be partially masked by the strong phenotypes induced by *cacn-1(RNAi)*. Thus, we reasoned that the short identity shared between *Y111B2A.25* and *cacn-1* dsRNA might induce strong sterility in the *sacy-1(tn1385*rf) genetic background but not in the wild type through weaker interference with *cacn-1*. To test this possibility, we systematically reduced the efficacy of the *cacn-1(RNAi)* response by serially diluting the *cacn-1(RNAi)*-inducing bacteria with bacteria containing the empty vector control (L4440). Consistent with the possibility that *Y111B2A.25(RNAi*) targets *cacn-1*, limiting the efficacy of the *cacn-1(RNAi)* response revealed specific enhancement of sterility, gamete degeneration, and embryonic lethality in the *sacy-1(tn1385)* genetic background (Table S3). Notably, the response to limited *cacn-1(RNAi)* exhibited by *sacy-1(tn1385)* animals was remarkably similar to their response to *Y111B2A.25(RNAi)* (Table S3). The human and Drosophila orthologs of CACN-1 have been identified as components of spliceosomal C complexes (Jurica *et al*. 2002; Bessonov *et al*. 2008, 2010; Herold *et al*. 2009; Fica *et al*. 2019). Like DDX41/Abstrakt, Cactin is recruited to the C complex of the spliceosome.

**Table 3.**
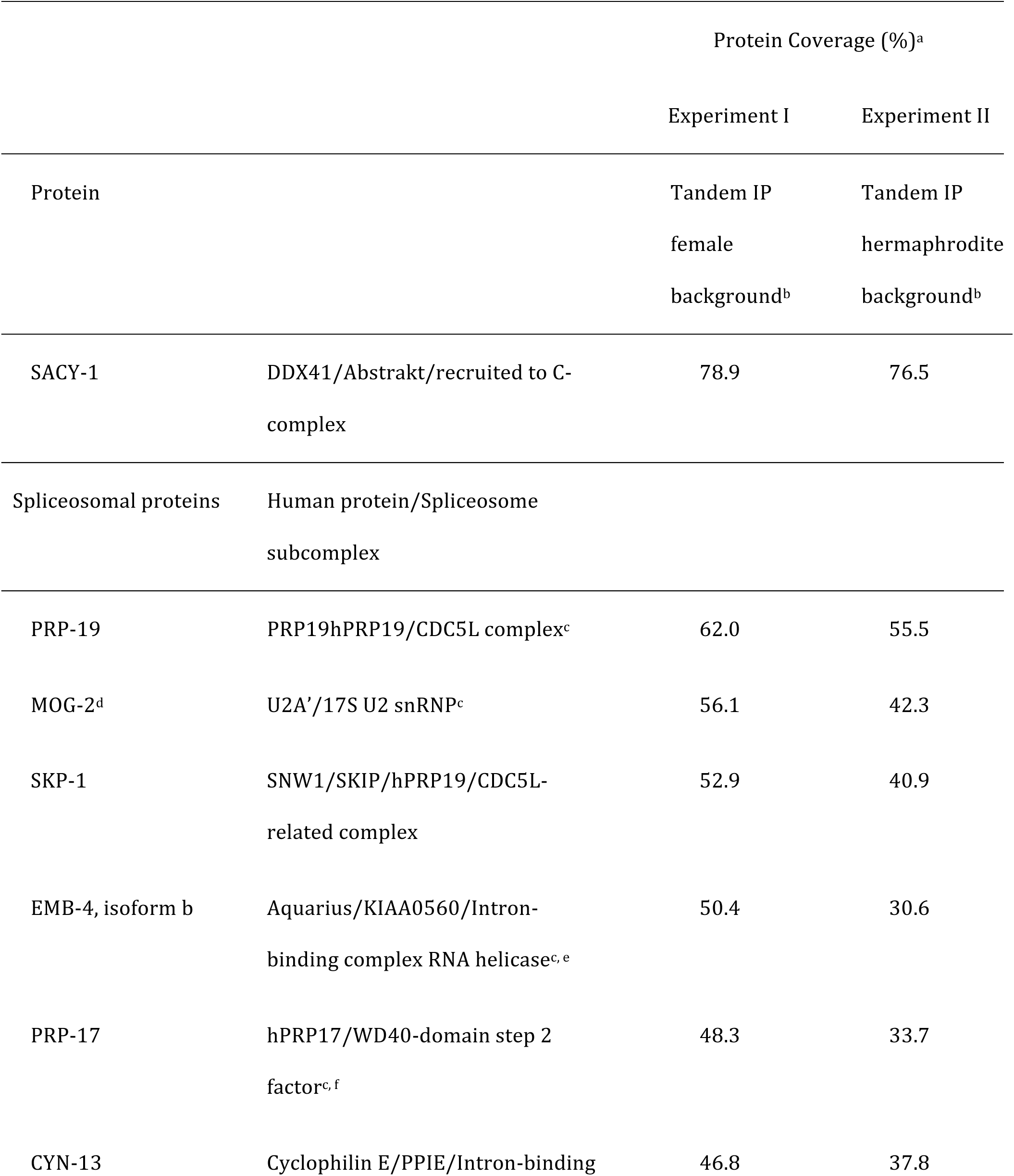
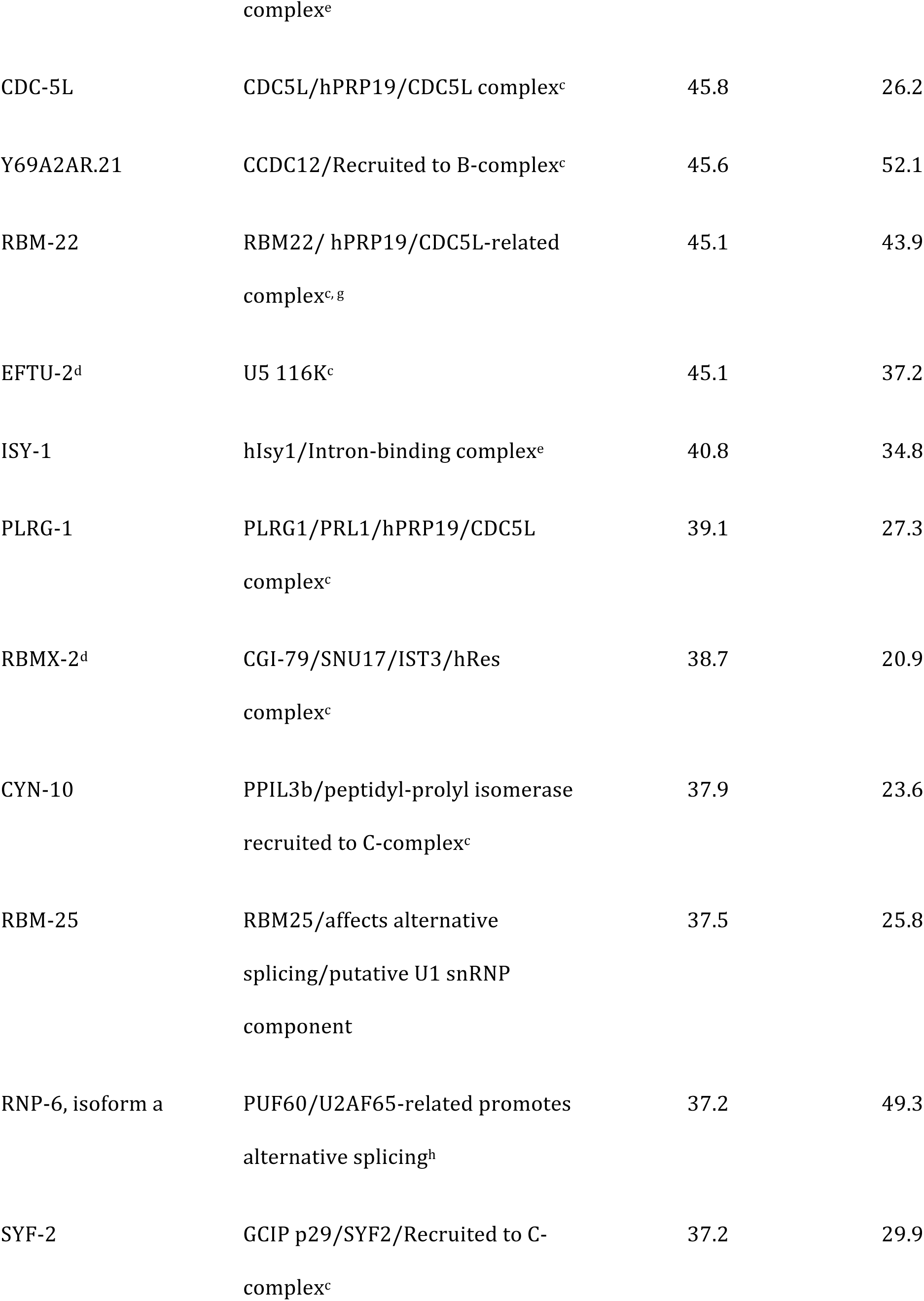
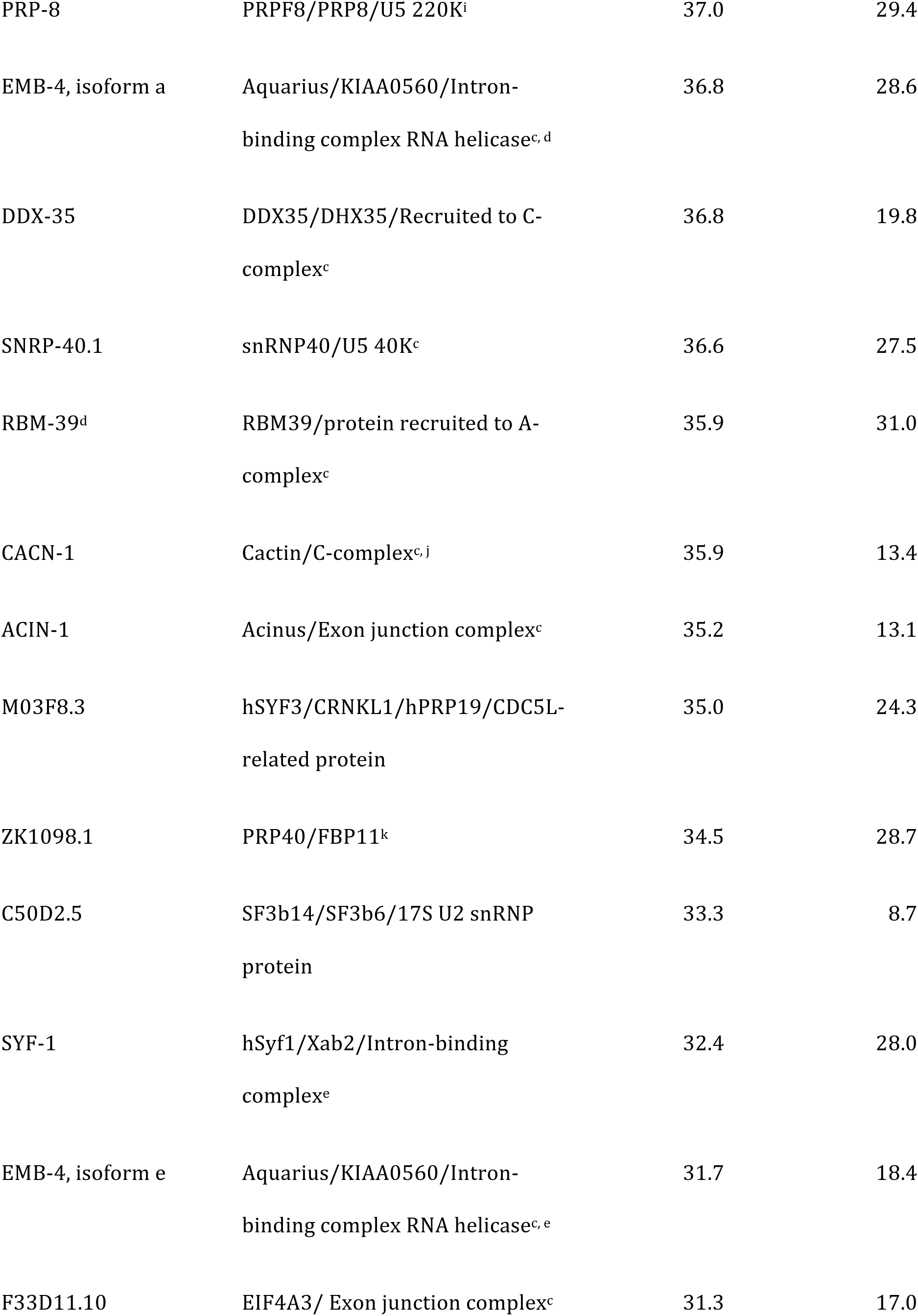
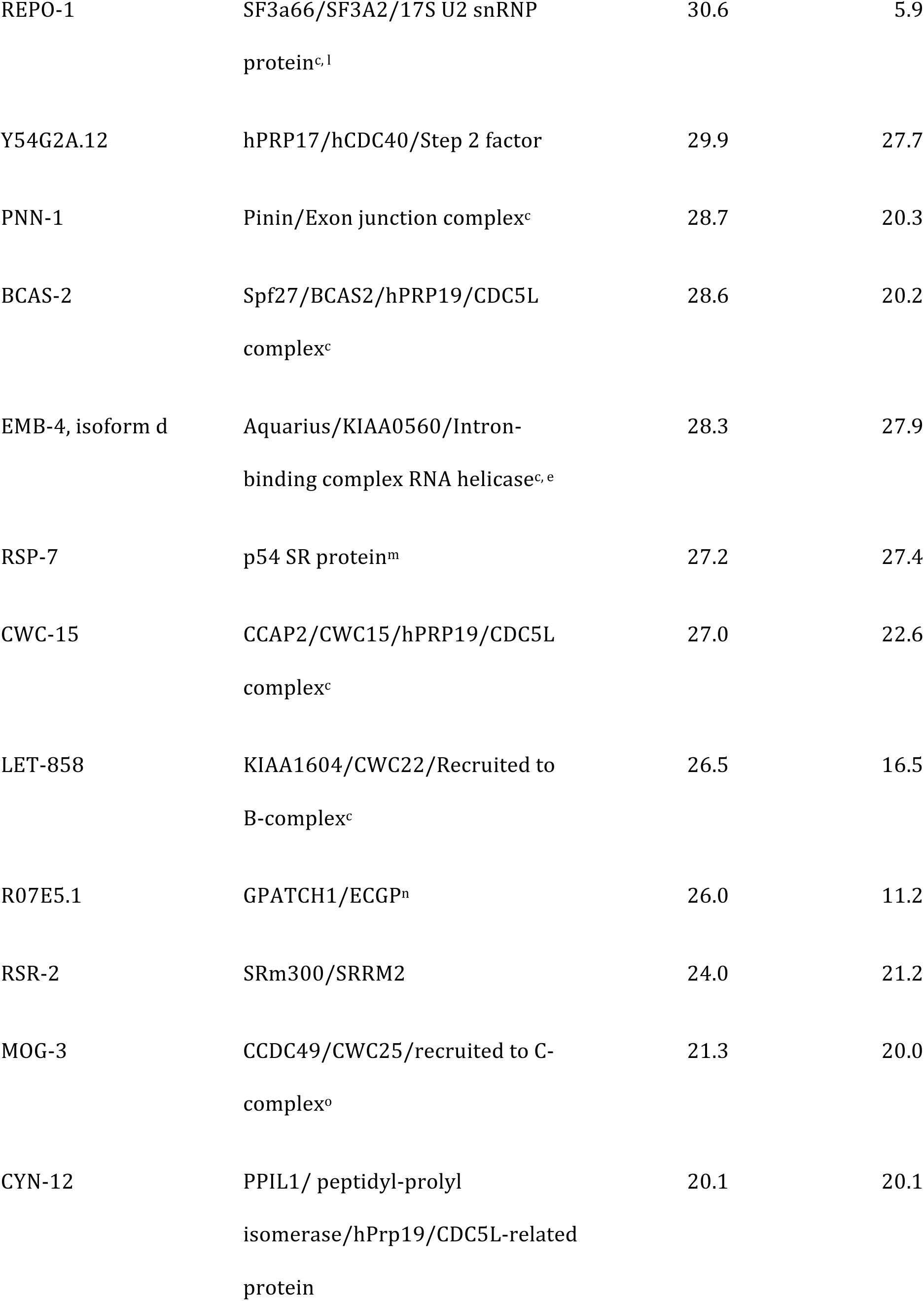
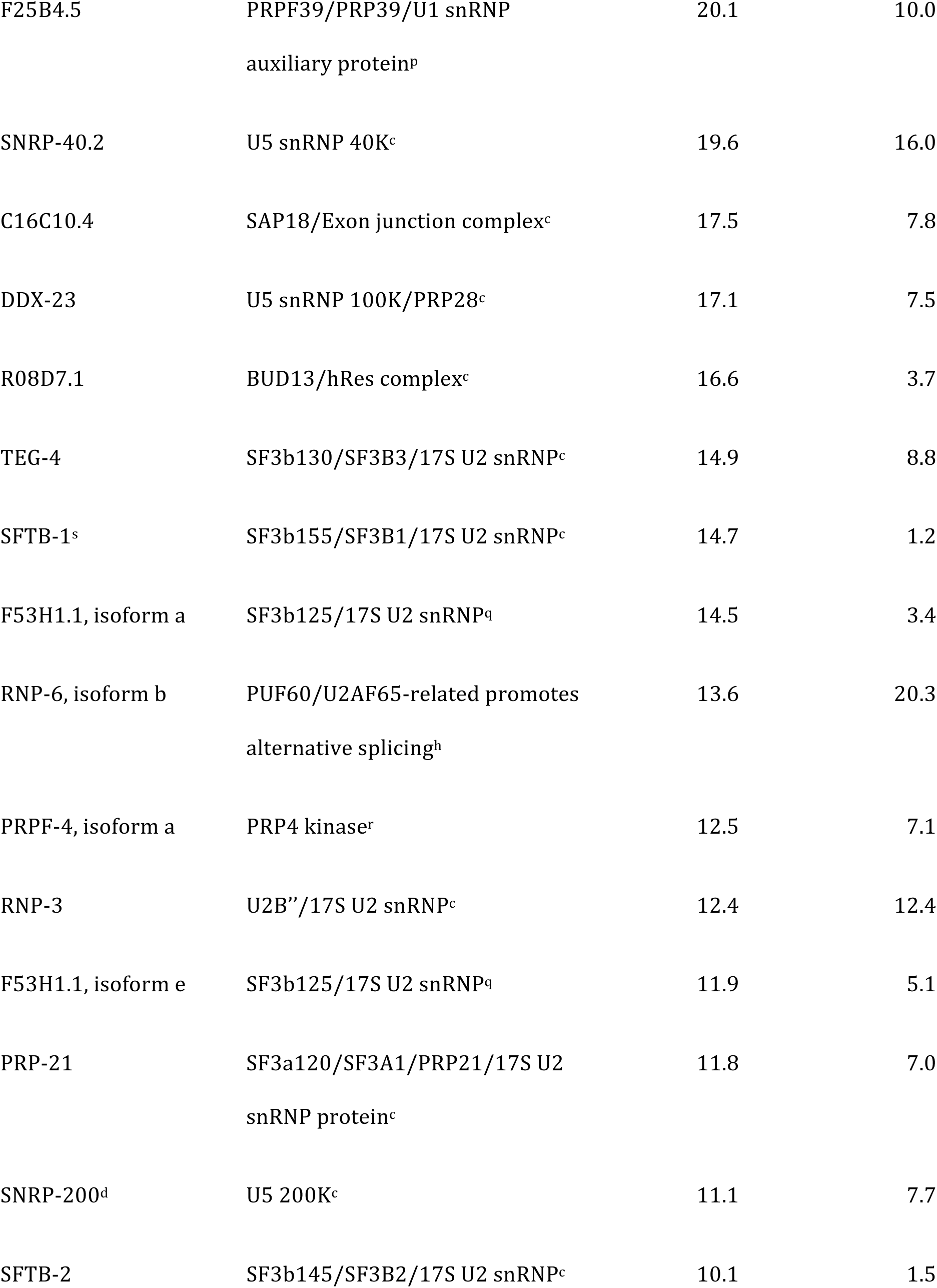
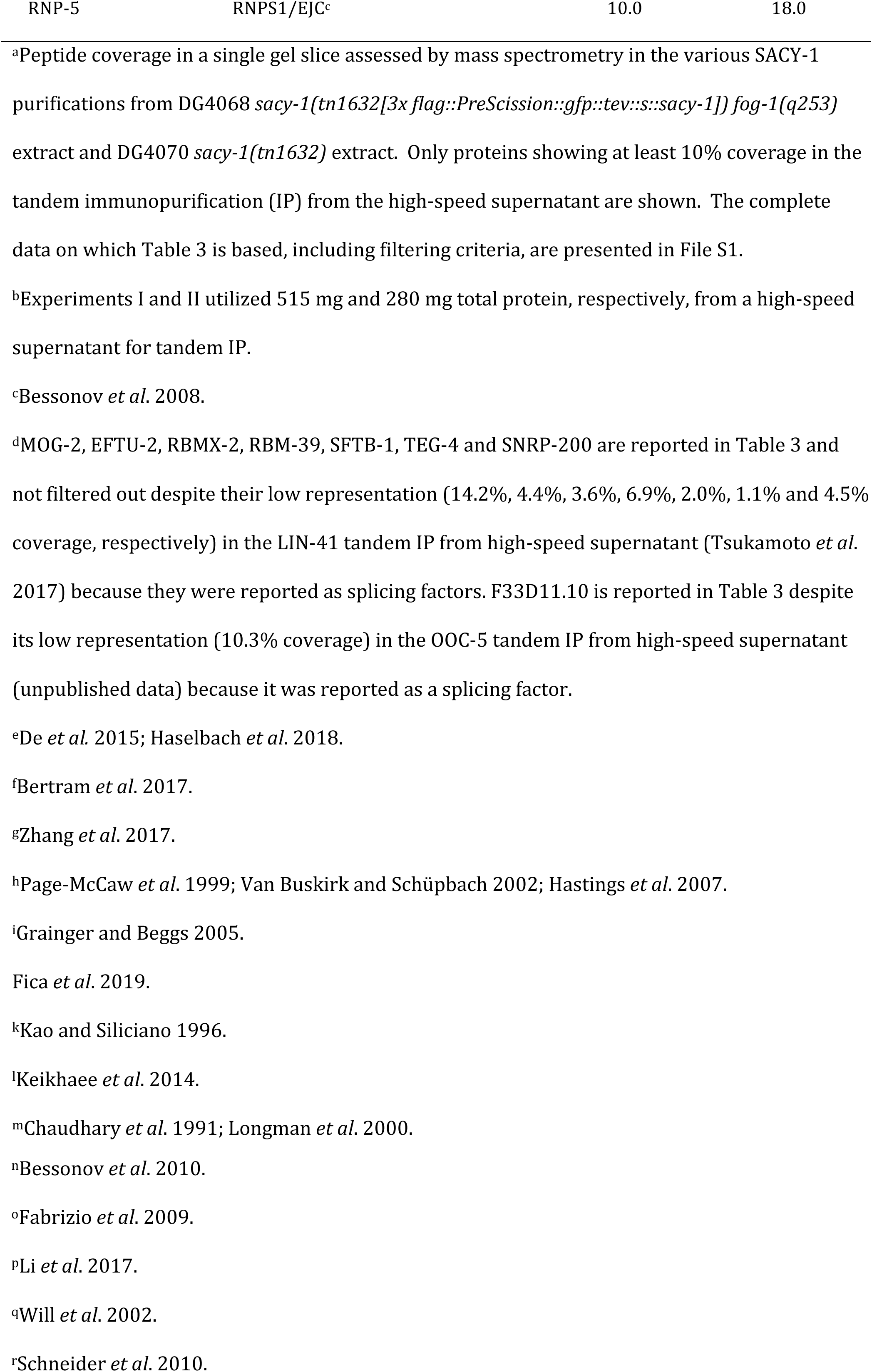
Spliceosomal proteins associated with SACY-1 using tandem affinity purification

In addition to *mog-2* and *Y111B2A.25*, we identified three different RNAi clones targeting the *emb-4* locus as strong enhancers of the *sacy-1(tn1385)* sterility and gamete degeneration phenotypes (Table 1; Figure S1). *emb-4* encodes a nuclear protein orthologous to human Aquarius/IBP160/KIAA0560/fSAP164, an intron-binding spliceosomal protein with a helicase-like domain (Sam *et al*. 1998; Jurica *et al*. 2002; Bessonov *et al*. 2008; Herold *et al*. 2009; De *et al*. 2015; Haselbach *et al*. 2018). To extend these RNAi results, we examined genetic interactions between *sacy-1* and *emb-4*, employing the *emb-4(sa44)* reduction-of-function allele. When combined with the *sacy-1(tm5503)* null allele, we observed enhancement of lethal vulval rupture and protruding vulva (Pvl) phenotypes in *sacy-1(tm5503); emb-4(sa44)* double mutants (Table 2A). We also observed enhancement of these phenotypes in *sacy-1(tn1385); emb-4(sa44)* double mutants, which were derived from *sacy-1(tn1385)/+; emb-4(sa44)* parents (Table 2A). Interestingly, the F1 progeny of fertile *sacy-1(tn1385); emb-4(sa44)* homozygous adults exclusively produced dead embryos or arrested L1-stage larvae, unlike each of the single mutants, which were highly fertile (Table 2B). Taken together these genetic interactions between *sacy-1* and three genes encoding spliceosomal proteins suggest that multiple *sacy-1* mutant phenotypes might result from compromised functions of the spliceosome.

### SACY-1 is a component of the C. elegans spliceosome

To characterize SACY-1-associated proteins, we conducted tandem affinity purifications using strains in which we used CRISPR-Cas9 genome editing to insert 3xFLAG and eGFP affinity tags at the SACY-1 N-terminus, separated by a PreScission protease recognition sequence (Figure S2). The resulting *sacy-1(tn1632[3x flag::PreScission::gfp::tev::s-tag::sacy-1])* strain was viable and fertile and exhibited no apparent germline or somatic defects. Although 3xFLAG::GFP::SACY-1 is expressed in all cells, it is particularly abundant in the female germline. Thus we conducted purifications from protein lysates prepared from adult animals in which the germline was feminized (experiment I) and also from adult hermaphrodites (experiment II). In both experiments, we found that 3xFLAG::GFP::SACY-1 associated with 55 proteins defined as spliceosomal proteins in other systems (Table 3; Jurica *et al*. 2002; Bessonov *et al*. 2008). The spliceosomal proteins identified in our genome-wide RNAi screen for *sacy-*1 enhancers (MOG-2, EMB-4, and CACN-1), were very well represented in our purifications (∼35–56% peptide coverage; Table 3). We also detected 9 additional spliceosomal proteins in the purification from the female but not the hermaphrodite genetic background (Table S4), but this might be a consequence of the fact that more protein extract was used in that experiment. We also detected 28 other proteins in our tandem affinity purifications (Table S5). Homologs of several of these factors have been implicated in the regulation of RNA splicing, including NRDE-2 (Jiao *et al*. 2019), CIR-1 (Maita *et al*. 2005; Kasturi *et al*. 2010), and CDK-12 (Rodrigues *et al*. 2012). These biochemical studies, taken together with the results from the genome-wide RNAi screen, suggest that both specific and pleiotropic defects conferred by *sacy-1* mutant alleles might result from spliceosomal defects.

**Table 4.** *sacy-1(tn1385rf)* enhances the *glp-1(ar202)* Tumorous phenotype

| Strain | Gonad arms containing mitotic undifferentiated germ cells in the proximal gonad arm<sup>a</sup> |
| --- | --- |
| *sacy-1(tn1385[G533R])* | 0 (n=256)<sup>b</sup> |
| *glp-1(ar202)* | 0.8 (n=364)<sup>b</sup> |
| *sacy-1(tn1385[G533R]); glp-1(ar202)* | 49.6 (n=415)<sup>b</sup> |
| *sacy-1(tn1887[R534H]); glp-1(ar202)* | 0 (n=62) |

<sup>a</sup>The percentage of young adult hermaphrodites were examined by DIC microscopy approximately 40 hours post-L4 at 15°C. The number of gonad arms scored is listed.
<sup>b</sup>In addition, dissected and fixed gonad from the same stage were stained for the phosphohistone H3(Ser10) M-phase marker. Of 25 *sacy-1(tn1385); glp-1(ar202)* gonads scored, 16 (64%) contained phosphohistone H3-positive undifferentiated germ cells in the proximal gonad arm. The average number of proximal phosphohistone H3-positive germ cells in the Tumorous gonads was 22 ± 12. None of the *sacy-1(tn1385)* (n=18) or *glp-1(ar202)* (n=31) dissected gonads examined contained phosphohistone H3-positive undifferentiated germ cells in the proximal gonad arm.

**Table 5.**
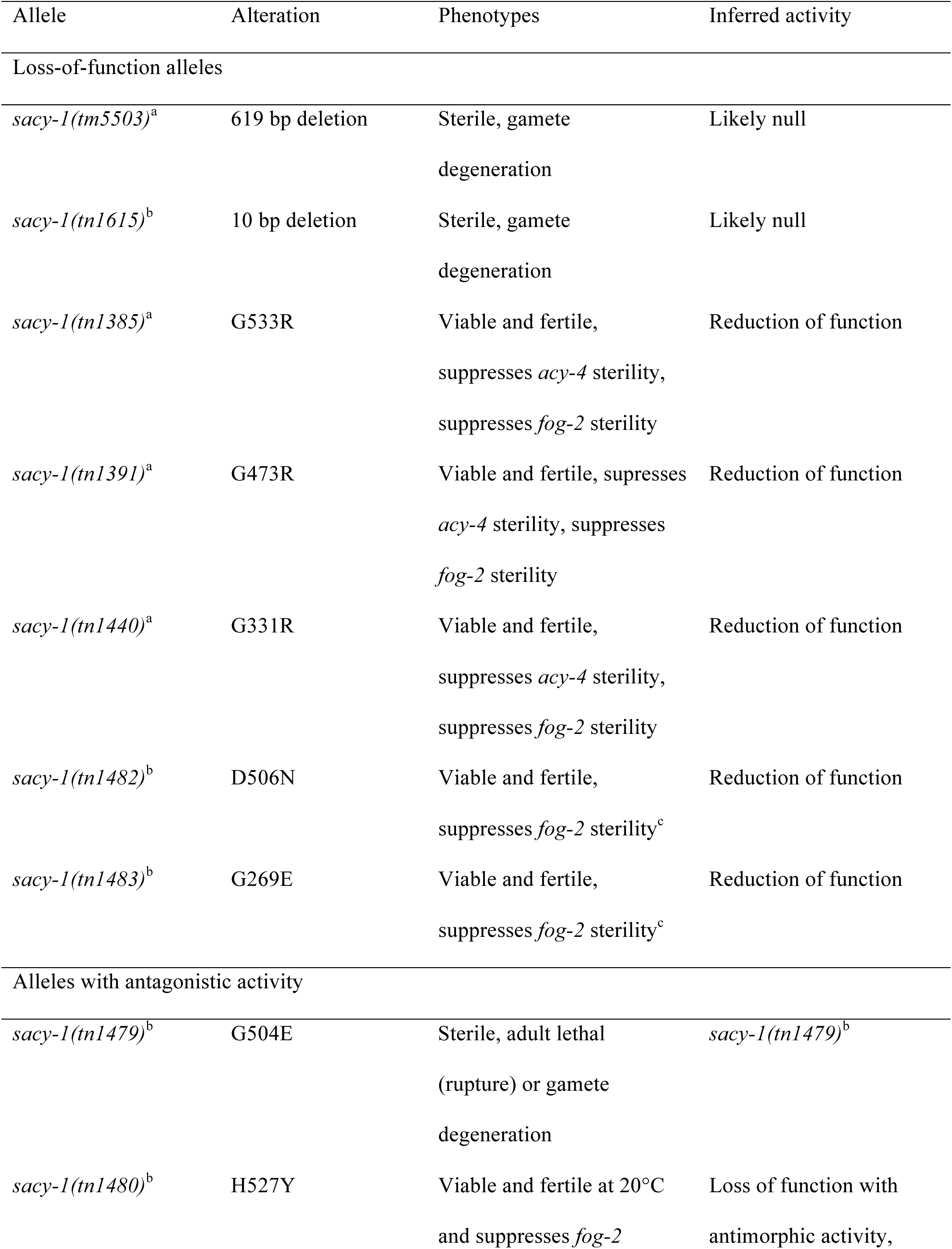
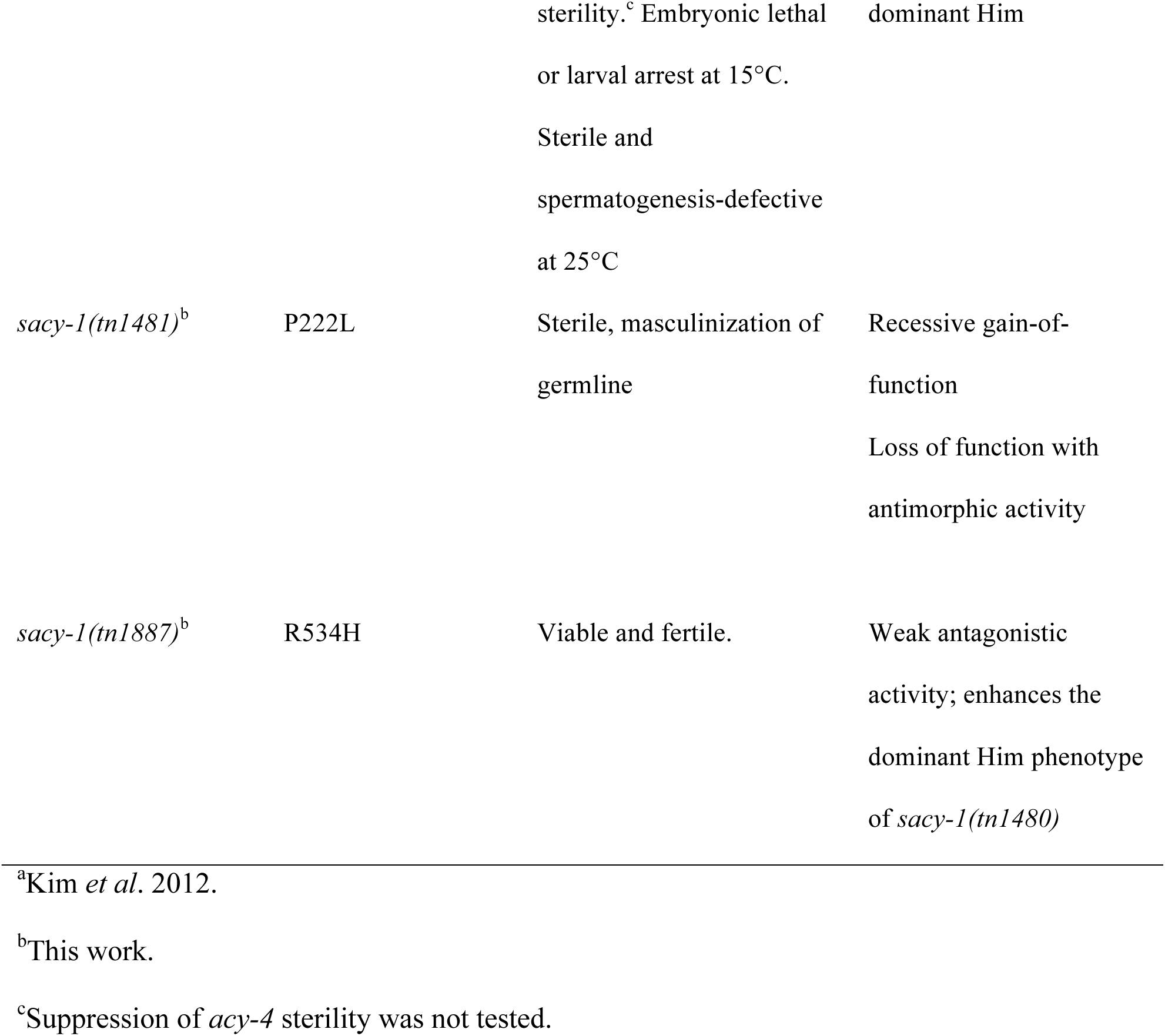
*sacy-1* alleles relevant to this study

### A sacy-1 reduction-of-function mutation enhances the Tumorous phenotype of a gain-of-function glp-1/Notch allele

Prior work showed that mutational or RNAi treatments affecting the function of multiple spliceosomal components could enhance weak gain-of-function (gf) mutations in *glp-1/Notch*, resulting in the ectopic proliferation of undifferentiated germ cells in the proximal gonad arm (Mantina *et al*. 2009; Kerins *et al*. 2010; Wang *et al*. 2012), which is referred to as a proximal proliferation or Tumorous phenotype. Thus, we constructed double mutants between the *sacy-1(tn1385)* mutation and the *glp-1(ar202*gf) mutation (Pepper *et al*. 2003). When analyzed in the young adult stage at 15°C (40 hrs post-L4), very few *glp-1(ar202*gf) adult hermaphrodites (∼0.8%) were observed to exhibit a proximal proliferation phenotype with undifferentiated germ cells in the proximal gonad arm (Table 4). By contrast, many *sacy-1(tn1385*rf*); glp-1(ar202*gf) adults (∼51%) exhibited a Tumorous phenotype (Table 4). This phenotype was not observed in *sacy-1(tn1385*rf) single mutants (Table 4). This result is consistent with the idea that the *sacy-1(tn1385*rf) mutation, though homozygous viable and fertile (brood size ∼350; Kim *et al*. 2012), compromises the function of the spliceosome, as assessed in a sensitized genetic background.

### Reduction-of-function sacy-1 mutations in C. elegans affect highly conserved residues in the DEAD-box and helicase domains

To better understand the functions and activities of the highly conserved SACY-1/DDX41 protein (Figure 1B), we conducted forward genetic screens for new *sacy-1* mutations, taking advantage of the fact that reductions of *sacy-1* function by mutation or RNAi can suppress the self-sterility of *fog-2* null mutations (Kim *et al*. 2012), which is caused by a failure to produce sperm (Schedl and Kimble 1988). Thus, we conducted a non-complementation screen for new mutations that enable fertility in trans to the *sacy-1(tn1385)* reduction-of-function allele in the *fog-2(oz40)* genetic background (Figure S3). In a screen of 15,577 haploid genomes, we isolated five new *sacy-1* missense alleles (*tn1479*–*tn1483*; Figure 1; see Table 5 for a list of all *sacy-1* alleles central to this work and their properties). All the *sacy-1* missense mutations isolated thus far in *C. elegans* alter highly conserved amino acids, and several of these mutations are nearby or in subdomains of the DEAD-box affected by DDX41 mutations found in human neoplasms (Figure 1B). The *sacy-1* missense alleles were modeled onto the crystal structures of DEADc and the HELICc domains of DDX41 (Schütz *et al*. 2010; Omura *et al*. 2016) and found likely to be surface accessible (Figure 1C), suggesting that the mutant alleles might interfere with the function of other protein components of the spliceosome. Consistent with this idea, the non-complementation screen resulted in the isolation of three novel *sacy-1* alleles (*tn1479*, *tn1480*, and *tn1481*), which appear to confer antagonistic activities, likely at the level of the spliceosome (see below).

**Figure 1.**
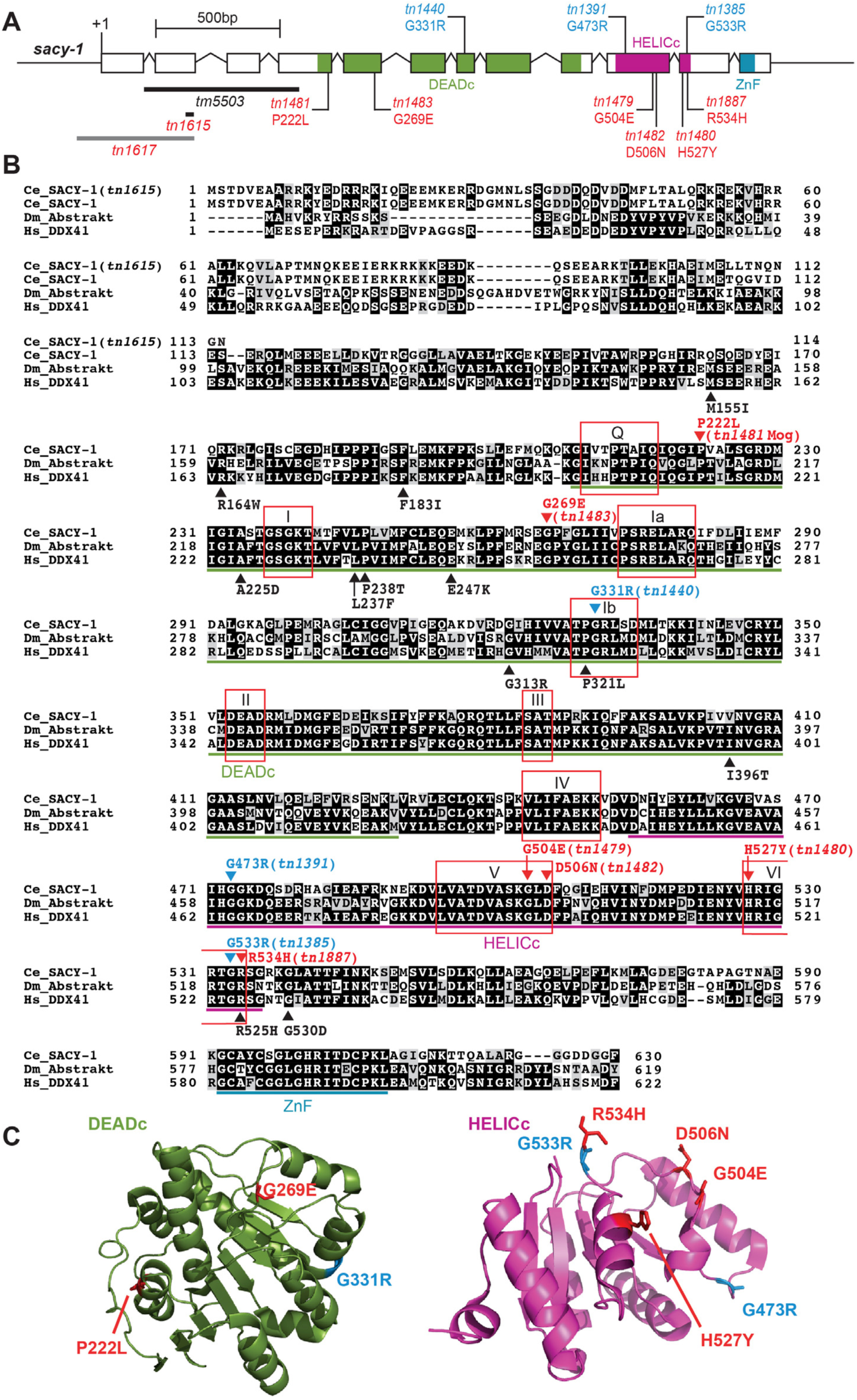
(A) The structure of *sacy-1*. Newly isolated mutations reported in this study are displayed in red font beneath the exons. The mutations in blue font shown above the exons were reported previously (Kim *et al*. 2012). The extent of two deletions, *tm5503* and *tn1615*, that result in *sacy-1* null mutations are shown with black bars. A third deletion, *tn1617*, which is a reduction-of-function mutation, is shown with a gray bar. (B) A protein sequence alignment of SACY-1 (NP_491962.1), Drosophila Abstract (NP_524220.1), and human DDX41 (NP_057306.2). Mutations isolated in *C. elegans* are shown above that sequence, whereas the human mutations associated with myelodysplastic syndromes are shown beneath the human sequence. Conserved domains [DEAD-box domain (DEADc), helicase domain (HELICc), and zinc finger domain (ZnF)] and motifs (Q, I Ia, Ib, III, IV, V, and VI) are indicated as described by Henn *et al*. (2012). (C) The locations of SACY-1 missense mutations are shown on structures of the DDX41 DEADc (Omura *et al*. 2016) and HELICc (Schütz *et al*. 2010) domains. The side chains of the amino acids in the human structure are labeled with amino acid numbering that corresponds to the SACY-1 missense mutations in this study.

Similar to the *sacy-1(tm5503)* deletion allele which removes exons 2 and 3 and a portion of exon 4 (Figure 1A; Kim *et al*. 2012), *sacy-1(tn1479)* homozygous hermaphrodites were sterile and displayed the gamete degeneration phenotype, but with reduced penetrance (Table 6; Figure S4). Consistent with the idea that *sacy-1(tm5503)* defines the null phenotype, an antibody specific to a portion of the DEAD-box domain downstream of the *tm5503* deletion (residues 411–578) fails to detect a protein product in extracts from *sacy-1(tm5503)* adults (Figure S2). To further define the *sacy-1* null phenotype, we used CRISPR/Cas9 genome editing to generate indels upstream of the DEAD-box-encoding regions by targeting Cas9 double-strand DNA breaks to exon 2 with an efficient sgRNA. We generated *sacy-1* indels in both wild-type as well as *lin-41(tn1541[gfp::lin-41])* hermaphrodites, the latter serving to provide a marker for oocyte development (Spike *et al*. 2014a,b). In these experiments, we generated 14 new *sacy-1* alleles (*tn1602*–*tn1612* and *tn1615*–*tn1617*). Of these, 13 displayed the gamete degeneration phenotype, again consistent with this representing the null phenotype. Not surprisingly, GFP::LIN-41 levels declined and the protein became undetectable as oocytes degenerated (D. Greenstein, unpublished results). One of the new candidate null alleles, *sacy-1(tn1615)*, was sequenced and found to result from a 10 bp deletion at the end of exon 2, which is predicted to introduce a stop codon prior to the DEAD-box domain, consistent with a null mutation (Figure 1B). Among the CRISPR-Cas9-induced alleles, *sacy-1(tn1617)* was exceptional in that it was homozygous viable and fertile, though slow growing, despite the fact that the deletion removes the initiation codon of *sacy-1* (Figure 1A). This exceptional allele might utilize an alternative start codon just prior to the DEAD-box domain, although this possibility was not explored. Since null mutations in *sacy-1* result in hermaphrodite sterility, there is the possibility that maternal *sacy-1(+)* activity contributes to the development of the germline and soma. Indeed, when the gamete degeneration phenotype is delayed through germline feminization, the mating of *sacy-1(tm5503)* null females to wild-type males produces embryos that arrest prior to morphogenesis (Kim *et al*. 2012).

**Table 6.** *sacy-1* mutant alleles with antagonistic activity

| Allele | Class | Gamete<br>Degeneration<sup>a</sup> | Vulval<br>Rupture<sup>a</sup> | Mog<sup>a,b</sup> | T<sup>c</sup> |
| --- | --- | --- | --- | --- | --- |
| *sacy-1(tm5503)*<br>(n=143) | Strong loss-of-function | 5.6 | 94.4 | 0 | 15 |
| *sacy-1(tm5503)*<br>(n=96) | Strong loss-of-function | 96.9 | 3.1 | 0 | 20 |
| *sacy-1(tm5503)*<br>(n=106) | Strong loss-of-function<br>(from *tm5503/+*) | 98.0 | 2.0 | 0 | 25 |
| *sacy-1(tm5503)<sup>d</sup>*<br>(n=231) | Strong loss-of-function<br>(from *tm5503/tn1480*) | 14.7 | 85.3 | 0 | 25 |
| *sacy-1(tn1615)*<br>(n=92) | Strong loss-of-function | 97.8 | 2.2 | 0 | 20 |
| *sacy-1(tn1479)*<br>(n=170) | Strong loss-of-function with<br>antagonistic activity | 11.2 | 88.8 | 0 | 20 |
| *sacy-1(tn1481)*<br>(n=125) | Recessive gain-of-function | 0 | 0 | 100 | 20 |

<sup>a</sup>The percentage of adult hermaphrodites exhibiting the reported phenotype is shown. Adults were scored 24 hr post-L4 at 20° or 25°C or 48 hr post-L4 at 15°C.
<sup>b</sup>*sacy-1(tn1481)* adult hermaphrodites produce large numbers of sperm but no oocytes and are sterile.
<sup>c</sup>Growth temperature in °C.
<sup>d</sup>The *sacy-1(tm5503)* progeny of *sacy-1(tm5503)/unc-13(e1091) sacy-1(tn1480)* hermaphrodites grown at 25°C.

One notable difference between *sacy-1(tn1479)* and the *sacy-1(tm5503)* null allele is that the majority of *sacy-1(tn1479*) adult hermaphrodites die by vulval rupture at 20°C (Table 6; Figure S4). This phenotype is only observed in a small minority of *sacy-1(tm5503)* hermaphrodites at 20°C but becomes the predominant phenotype at 15°C (Table 6). This observation suggests that *sacy-1(tn1479)* is more severe than a null allele, as might happen if the SACY-1[G504E] product is nonfunctional but also antagonizes a protein complex containing SACY-1, likely the spliceosome.

Unlike the *sacy-1(tm5503)* null allele, which is sterile, three of the newly isolated mutations *(tn1480*, *tn1482*, and *tn1483*) are self-fertile as homozygotes at 20°C similar to the *sacy-1(tn1385*rf) mutant. However, *sacy-1(tn1480)* and *sacy-1(tn1483)* homozygotes exhibited additional defects; both appeared to grow more slowly than the wild type at this temperature. In addition, *sacy-1(tn1483)* adult hermaphrodites had smaller gonad arms, suggesting effects on germline proliferation (T. Tsukamoto and D. Greenstein, unpublished results). Finally, *sacy-1(tn1480)* confers temperature-dependent gain-of-function pleiotropic defects that will be described below.

### Novel sacy-1 mutant alleles appear to antagonize essential functions of the spliceosome

#### A recessive gain-of-function sacy-1 mutation masculinizes the hermaphrodite germline

In the non-complementation screen for *sacy-1* mutant alleles, we isolated a novel *sacy-1* allele, *tn1481*, with a masculinization of germline (Mog) phenotype (Table 6; Figure 2). All *sacy-1(tn1481)* homozygous hermaphrodites produce excess numbers of sperm but no oocytes (n=125). Staining of dissected gonads from adults showed that whereas all wild-type hermaphrodite gonad arms examined (n=64) expressed both the major sperm protein (MSP) and the RME-2 oocyte yolk receptor, all *sacy*-*1(tn1481)* gonad arms (n=178) expressed only MSP but not RME-2 (Figure 2). In our non-complementation screen (Figure S3), we also isolated a *gld-1* Mog allele, *tn1478*, as a dominant suppressor of *fog-2(oz40)*. *gld-1(tn1478)* results from the same G248R amino acid substitution reported for the *gld-1(q93)* Mog allele (Francis *et al*. 1995a,b; Jones and Schedl 1995). Thus it was important to determine whether the *sacy-1(tn1481*[P222L]) mutation was the cause of the Mog phenotype. This was ascertained by crossing a GFP::SACY-1 transgene (*tnEx159*) into the *sacy*-*1(tn1481)* genetic background. We found that all *sacy-1(tn1481); tnEx159 [gfp::sacy-1 +unc-119(+)]* hermaphrodites (n=30) produced oocytes and sperm and were self-fertile. This result established that the P222L mutation in SACY-1 causes the Mog phenotype.

**Figure 2.**
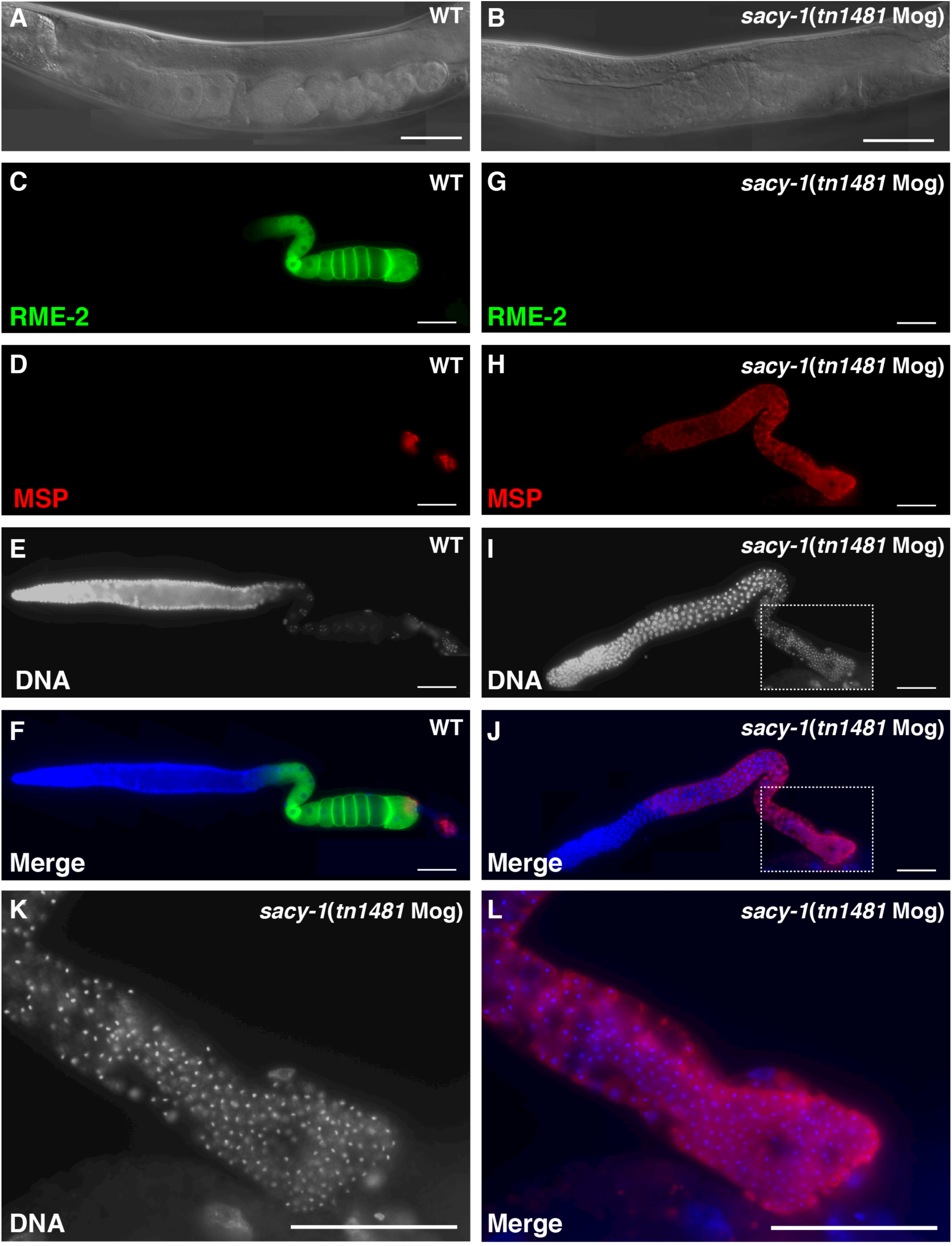
*sacy-1(tn1481)* adult hermaphrodites exhibit a masculinization of germline (Mog) sterile phenotype. DIC images of wild-type (A) and *sacy-1(tn1481)* (B) adult hermaphrodites. The wild-type animal contains oocytes and sperm and produces embryos but the *sacy-1(tn1481*) animal only produces sperm (arrow). (C–L) Dissected gonads stained for the RME-2 yolk receptor (C, G), the major sperm protein (D, H), or DNA (E, I, and K). Merged images are also shown (F, J, and L). The *sacy-1(tn1481)* mutant overproduces sperm to the exclusion of oocytes and is sterile. This phenotype is completely penetrant. Bars, 50 μm.

The suppression of *fog-2* sterility by reduction-of-function *sacy-1* mutations is consistent with *sacy-1(+)* possessing a function that promotes the oocyte fate; this function is non-essential, however, because the strongest loss-of-function *sacy-1* alleles are able to produce oocytes, which nevertheless undergo necrotic degeneration. Thus, the *sacy-1(tn1481)* Mog phenotype suggests this mutant allele, although recessive, possesses an activity antagonistic to this oocyte-promoting function in the sperm-to-oocyte switch. To genetically characterize *sacy-1(tn1481)* further, we analyzed the phenotype of *sacy-1(tn1481)/sacy-1(tm5503* null) heterozygotes. Whereas all *sacy-1(tn1481)* homozygotes (n=50) displayed a Mog phenotype, all *sacy-1(tn1481)/sacy-1(tm5503)* heterozygotes (n=48) produced both oocytes and sperm and were self fertile. This result suggests that the *sacy-1(tn1481)* Mog phenotype is dosage sensitive and that the mutant allele appears to be a recessive gain-of-function mutation that might antagonize proteins that normally function with SACY-1, likely other components of the spliceosome as discussed below.

In *C. elegans*, a genetic hierarchy controls germline sex determination (Figure 3). The failure of *sacy-1(RNAi)* to suppress the sterility of the dominant strongly feminizing *tra-2(e2020)* mutation, which deletes GLD-1 binding sites within the *tra-2 3’-UTR*, was interpreted in the context of a model in which *sacy-1(+)* promotes the oocyte fate in opposition to *fog-2* and *gld-1* at the level of *tra-2* (Figure 3; Kim *et al*. 2012). Because the evaluation of potential interactions between *sacy-1* and *tra-2* relied on *sacy-1(RNAi)*, there was the concern that this treatment reduced but did not eliminate the function of *sacy-1*. Thus, we reevaluated the interaction between *tra-2* and *sacy-1* genetically. In the first approach, we combined the *sacy-1(tm5503)* null allele with *tra-2(e2020)*. We analyzed the sexual fate of the germline by staining dissected gonads from adult animals with oocyte (RME-2) and sperm (MSP) markers. Whereas all wild-type gonad arms examined (n=30) expressed RME-2 and MSP, all gonad arms of *sacy-1(tm5503)*; *tra-2(e2020)* animals (n=26) expressed only RME-2 and not MSP (Figure 4). This result is consistent with the model in which *sacy-1* promotes the oocyte fate by promoting the function of *tra-2*. Although the germlines of *sacy-1(tm5503)*; *tra-2(e2020)* adults were feminized, oocytes underwent meiotic maturation constitutively, consistent with the finding that *sacy-1* is a negative regulator of meiotic maturation (Kim *et al*. 2012). *sacy-1(tm5503)*; *tra-2(e2020)* animals did however exhibit a highly penetrant ovulation defect, which caused endomitotic oocytes to accumulate in the gonad arm (Emo phenotype; Figure 4).

**Figure 3.**
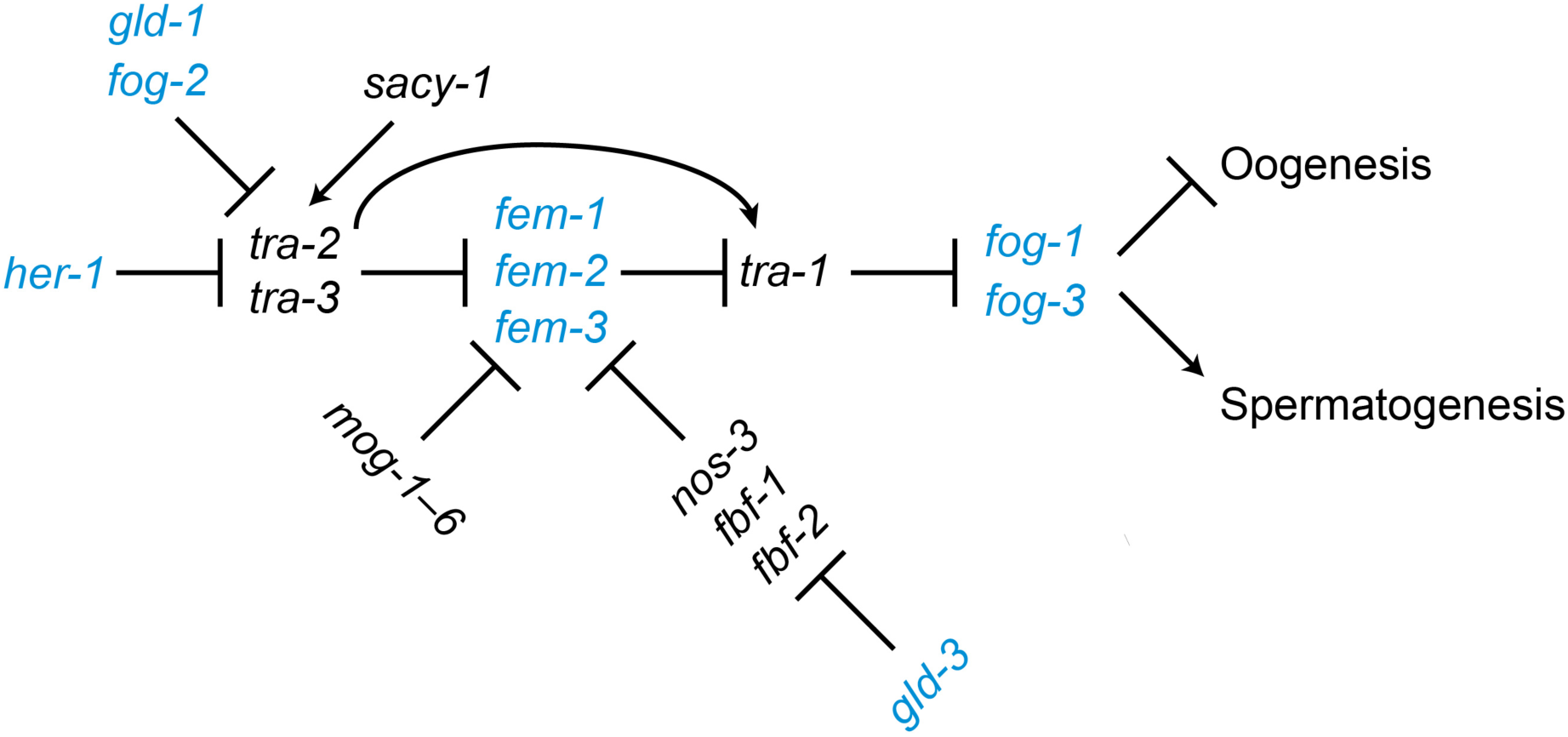
The *C. elegans* germline sex determination pathway. Genes promoting the male and female fate are shown in blue and black, respectively. *sacy-1* promotes the oocyte fate antagonistically to *fog-2*, which promotes spermatogenesis.

**Figure 4.**
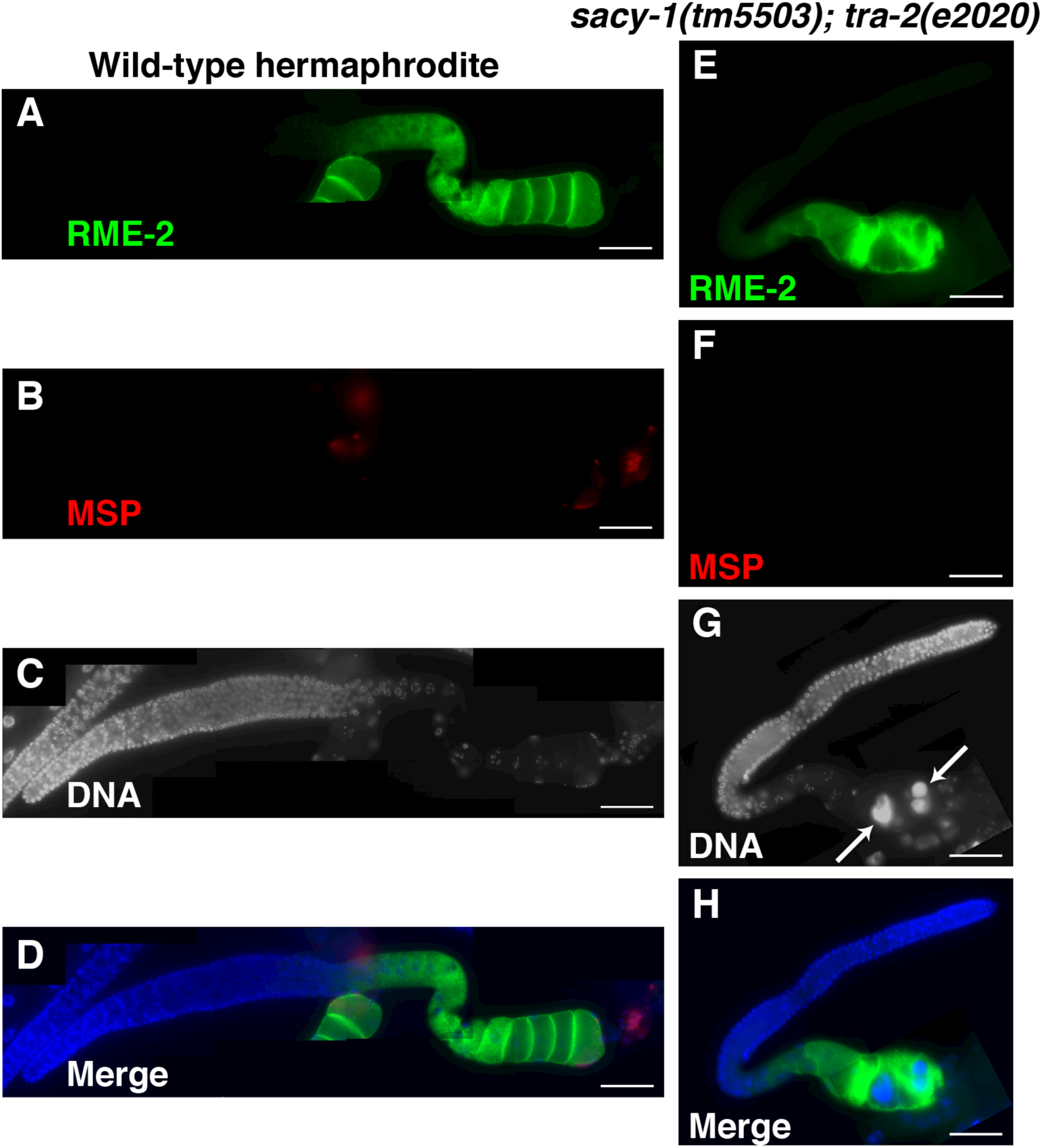
Genetic interactions between *sacy-1* and *tra-2* in germline sex determination were analyzed by combining the strong germline-feminizing dominant *tra-2(e2020)* mutation with the *sacy-1(tm5503)* null mutation. Dissected gonads of wild-type hermaphrodites (A–D) and *sacy-1(tm5503); tra-2(e2020)* females were analyzed by staining for the oocyte RME-2 yolk receptor (A and E) and the major sperm protein (B and F). DNA was detected with DAPI (C and G). Merged images are also shown (D and H). Note gonads from *sacy-1(tm5503); tra-2(e2020)* females do not express MSP and frequently contain endomitotic oocytes (arrows). Bars, 50 μm.

To extend these observations, we examined germline sexual fates in dissected gonads from *sacy-1(tn1481*Mog*); tra-2(e2020)* adults. Whereas all wild-type gonad arms examined (n=21) expressed MSP and contained sperm, none of the *sacy-1(tn1481Mog); tra-2(e2020)* gonad arms (n=37) expressed MSP or contained sperm. We noted that the Emo phenotype was less penetrant in *sacy-1(tn1481Mog); tra-2(e2020)* gonad arms (46% penetrance). These results are consistent with the possibility that *sacy-1(+)* promotes the oocyte fate by promoting the function of *tra-2* in the germline and suggests that *sacy-1(tn1481*Mog) may interfere with this function.

Interestingly, recessive loss-of-function mutations in six genes, *mog-1–6*, cause a Mog phenotype and encode spliceosomal components (Graham and Kimble 1993; Graham *et al*. 1993; Puoti and Kimble 1999, 2000; Belfiore *et al*. 2004; Zanetti *et al*. 2011). Mutation and RNAi depletion of many splicing factors have been observed to result in a Mog phenotype, suggesting that the germline sex determination process is particularly sensitive to disruptions in RNA splicing (Mantina *et al*. 2009; Kerins *et al*. 2010; Wang *et al*. 2012; Novak *et al*. 2015). Prior studies focusing on the *C. elegans* soma were interpreted in the context of a model in which *mog-1–mog-6* might function at the level of *fem-3* through 3’UTR-dependent translational regulation (Gallegos *et al*. 1998; Figure 3); however, the experiments in that study did not address the regulation of *fem-3* in the germline. We previously showed that *sacy-1(tm5503); fem-3(e1996)* adult XX animals had feminized germlines (Kim *et al*. 2012). To examine the genetic relationship between *sacy-1* and *fem-3* further, we generated *sacy*-*1(tn1481Mog); fem-3(e1996)* double mutants. We observed that 92% (n=23) of *sacy-1(tn1481Mog); fem-3(e1996)* animals were feminized. Mating of *sacy-1(tn1481Mog); fem-3(e1996)* females (n=29) to wild-type males resulted in the production of embryos that failed to hatch (n=4140, 99.8%) or arrested as larvae (n=7, 0.2%). This result indicates that two copies of *sacy-1(tn1481)* in the maternal germline, but not one [e.g., *sacy-1(tn1481)/sacy-1(tm5503)* heterozygotes are fertile] are incompatible with embryonic development. We found that 8% (n=2) of *sacy-1(tn1481*Mog*); fem-3(e1996)* animals (n=25) produced oocytes and sperm and a few dead embryos (one of these animals produced sperm in one gonad arm but not the other). This result suggests that the *sacy-1(tn1481)* mutation can promote sperm development independent of zygotic *fem-3(+)* activity, consistent the genetic epistasis pathway (Figure 3; Zanetti and Puoti 2013). We did observe that a reduction in *fem-3* dosage could suppress the *sacy-1(tn1481)* Mog phenotype (n=60). Specifically, whereas 65% (n=39) of *sacy-1(tn1481); fem-3(e1996)/+* animals were Mog, exclusively producing sperm, 33% (n=20) produced sperm and oocytes, and one animal (2%) was feminized. Since SACY-1 genetically and biochemically interacts with components of the spliceosome, we suggest that the *sacy-1(tn1481)* mutation antagonizes functions of the spliceosome needed for germline sex determination and oogenesis.

#### The gain-of-function sacy-1(tn1480) allele confers multiple pleiotropic phenotypes in a temperature-dependent manner

Interestingly, *sacy-1(tn1480)* displayed both cold-sensitive (15°C) and temperature-sensitive (25°C) defects. At 15°C, *sacy-1(tn1480)* homozygotes (n=55), which were the offspring of heterozygous parents grown at 15°C, laid dead eggs and produced arrested larvae (91%) or produced very few progeny (9%). At 25°C, *sacy-1(tn1480)* hermaphrodites were sterile and produced abnormal and fertilization-defective sperm (Figure 5). *sacy-1(tn1480)* hermaphrodite sterility is rescued by mating with wild-type males at 25°C. In addition to producing oocytes, some *sacy-1(tn1480)* adult hermaphrodites continued to produce sperm, as swollen germ cells specified in the male fate were detected distal to the loop region in 42% of gonad arms examined (Figure 5; n=12). Thus, in addition to the other phenotypes it confers, the *sacy-1(tn1480)* mutation perturbs the sperm-to-oocyte switch at 25°C. We also observed that *sacy-1(tn1480)* conferred a dominant high-incidence of males (Him) phenotype; *sacy-1(tn1480)/+* heterozygous hermaphrodites produced 1.6% male progeny (n=3759), as compared to 0.1% for the wild-type control (n=6044; p<0.01, Fisher’s exact test). Genetic mapping showed that the temperature-dependent pleiotropic defects were inseparable from the *sacy-1(tn1480)* mutation (see Materials and Methods). Like *sacy-1(tn1481)*, the defects conferred by *sacy-1(tn1480)* were dosage sensitive. At 25°C, nearly all *sacy-1(tn1480)/sacy-1(tm5503)* heterozygotes (99%, n=88) were fertile (the average brood size was 132 ± 64, n=34); however, the majority of *sacy-1(tm5503)* homozygotes (85.3%, n=231) they produced burst as adults (Table 6). This result suggests that maternal *sacy-1(tn1480)* activity can antagonize the spliceosome in the absence of zygotic *sacy-1(+)* function. By these genetic criteria, *sacy-1(tn1480)* exhibits recessive and weakly dominant gain-of-function properties, depending on the phenotype examined.

**Figure 5.**
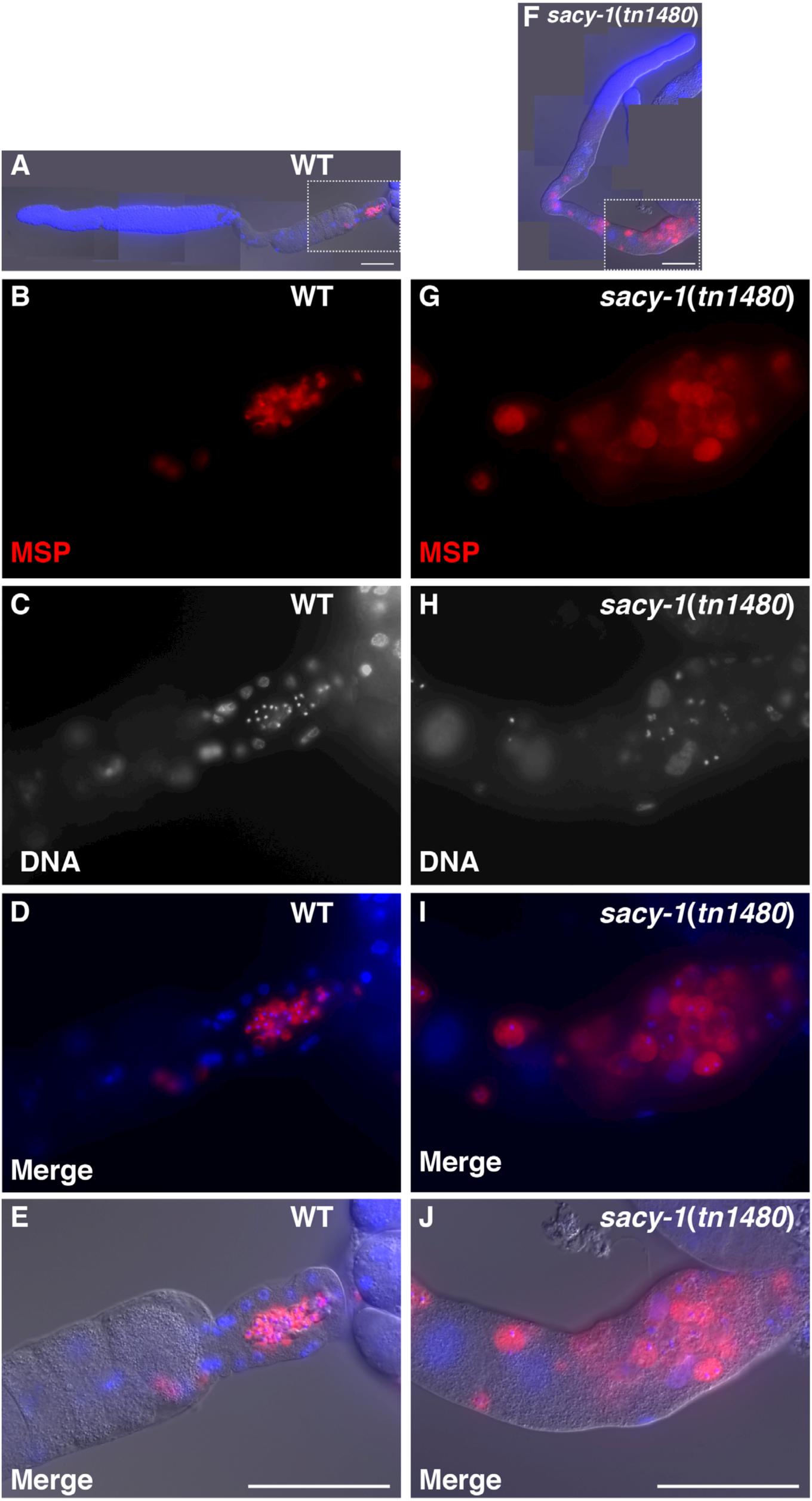
Sperm-defective phenotype of *sacy-1(tn1480)* at 25°C. Dissected gonads of wild-type (A–E) and *sacy-1(tn1480)* (F–J) stained with anti-MSP antibodies (B, G) and DAPI to detect DNA (C, H). Merged images are also shown (A, D, E, F, I, and J). At 25°C, *sacy-1(tn1480)* hermaphrodites produce swollen and abnormal sperm, which are incapable for fertilization. Note that defective sperm are also found near the bend region in *sacy-1(tn1480)* adults (F) indicating that there is a defect in the sperm-to-oocyte switch. Bars, 50 μm.

### The oncogenic DDX41 R525H mutation confers weak antagonistic activity in C. elegans

The R525H mutation in DDX41 has been reported in myeloid leukemias both as newly arising somatic mutations specific to the neoplastic cells, as well as inherited germline mutations (Polprasert *et al*. 2015; Lewinsohn *et al*. 2016; Sébert *et al*. 2019). Thus, it was of interest to examine the impact of the analogous mutation (R534H) on *sacy-1* function. Interestingly, a substitution at the adjacent amino acid (G533R) results in the *sacy-1(tn1385)* reduction-of-function mutation (Figure 1). Thus, we used genome editing to introduce the R534H mutation in the *C. elegans* genome (see Materials and Methods). By several criteria, *sacy-1(tn1887*[R534H]) homozygotes were indistinguishable from the wild type. Neither did *sacy-1(tn1887)* suppress *acy-4* sterility (n=140; *sacy-1(tn1887); acy-4(ok1806)* brood size was 1.5 ± 2.3) nor did it suppress *fog-2* sterility (n=48). All unmated *sacy-1(tn1887); fog-2(oz40)* gonad arms examined (n=96) exhibited stacked oocytes, which indicates that the *sacy-1(tn1887)* mutation does not derepress meiotic maturation in the absence of sperm, like strong reduction-of-function mutations do. Further, the *sacy-1(tn1887*[R534H]) did not enhance the Tumorous phenotype of a gain-of-function *glp-1/Notch* allele (Table 4). The brood size of *sacy-1(tn1887)* (204 ± 37; n=20) at 25°C was indistinguishable from that of the wild type (202 ± 62, n=30; p>0.8, two-sample Z-test). Also, the incidence of males (0.2%, n=4087) was similar to that observed in the wild type (0.1%, n= 6044). When placed in trans to the *sacy-1(tm5503)* null mutation, *sacy-1(tn1887)/sacy-1(tm5503)* heterozygotes were found to be fertile at all temperatures examined (15°, 20°, and 25°C (n=104)). However, we did observe that *sacy-1(tn1887)* significantly enhanced the dominant Him phenotype of *sacy-1(tn1480)* (p<0.05, Fisher’s exact test); *sacy-1(tn1887)/sacy-1(tn1480)* heterozygotes produced 5.7% males at 25°C (n=1483) as compared to 1.6% for the *+/sacy-1(tn1480)* control (n=3759). The brood size of *sacy-1(tn1887)/sacy-1(tn1480)* heterozygotes at 25°C (74 ± 65, n=20) was also significantly lower than that of the *+/sacy-1(tn1480)* control (188 ± 68, n=20; p<0.001, two-sample Z-test). Because, *sacy-1(tn1480)/sacy-1(tm5503)* heterozygotes do not exhibit an enhanced Him phenotype at 25°C (0.7% males, n=4493), the *sacy-1(tn1887*[R534H]) mutation appears to possess a weak antagonistic activity.

### Impact of sacy-1 on the transcriptome

#### Depletion of SACY-1 using the auxin-inducible degradation system

RNA sequencing studies using human patient samples suggested a role for DDX41 in splice site selection for a small number of human genes (Polprasert *et al*. 2015). Thus, we sought to address the impact of SACY-1 on the transcriptome by exploiting the power of the *C. elegans* system for transcriptomics under genotypically and experimentally well-controlled conditions. We chose to use the auxin-inducible degradation system (Zhang *et al*. 2015) to acutely deplete SACY-1 in the adult stage and thus avoid indirect impacts on the transcriptome arising as a developmental consequence of strong loss-of-function phenotypes (e.g., germline degeneration and cell fate changes). Because the genetic analysis established requirements for *sacy-1(+)* function in both the germline and soma, we used strains bearing germline (CA1352 *ieSi64*) or somatically expressed (CA1200 *ieSi57*) TIR1 F-box proteins to deplete AID::GFP::SACY-1 in each tissue individually (Figure S5 and Figure S6). Depletion of AID::GFP::SACY-1 in the germline starting at approximately the L3 stage phenocopied the gamete degeneration phenotype in a small proportion of the animals (3 of 270; Figure S5). This result is consistent with genetic mosaic analysis showing that the gamete degeneration phenotype is cell autonomous (Kim *et al*. 2012), and it also highlights the difficulty of recapitulating null phenotypes through auxin-inducible degradation. When AID::GFP::SACY-1 is depleted in the germline starting at the L4 stage, many of their F1 progeny arrest as embryos or larvae, consistent with the idea that maternally contributed *sacy-1(+)* activity is essential. Animals that escape the lethality and progress to adulthood often display the germline degeneration phenotype (40%; n=139). To deplete AID::GFP::SACY-1 in the soma, we placed L4 larvae on media containing 2 mM auxin. The resulting F1 progeny grew very slowly, taking approximately 4–6 days to reach adulthood (instead of 2.5 days) and were sterile (Figure S5). Taken together, depletion of SACY-1 using the auxin-inducible degradation system resulted in a reduction-of-function condition less severe than the null phenotype but more severe than conferred by the reduction-of-function missense alleles (Table 5).

For analysis of transcriptomes, we grew adult hermaphrodites to the young adult stage on normal growth media and then transferred them to media containing 2 mM auxin for 24 hours before preparing total RNA for RNA sequencing. Examination of the worms showed that AID::GFP::SACY-1 was depleted from the targeted tissues (Figure S6). Total RNA was prepared from each of three biological replicates and their respective controls, which were the parent strains expressing TIR1 in the germline (*ieSi64*) or soma (*ieSi57*) also treated with auxin. Poly(A+) mRNA was sequenced using 150 bp paired-end reads and the sequencing reads were aligned to the genome. Principal component analysis (PCA) revealed that the biological replicates clustered together (Figure 6A), which is indicative of experimental reproducibility. However, PCA indicated that the control strains for the germline (CA1352) and soma (CA1200) depleted samples did not cluster together, which indicates that under these conditions the two strain backgrounds exhibit marked differences in their transcriptomes (Figure 6A), a finding that was further confirmed with a more granular assessment of mRNA expression level differences of individual genes (Figure S7A). Thus, in our analysis we compared the germline- and soma-depleted SACY-1 transcriptomes only to their respective controls.

**Figure 6.**
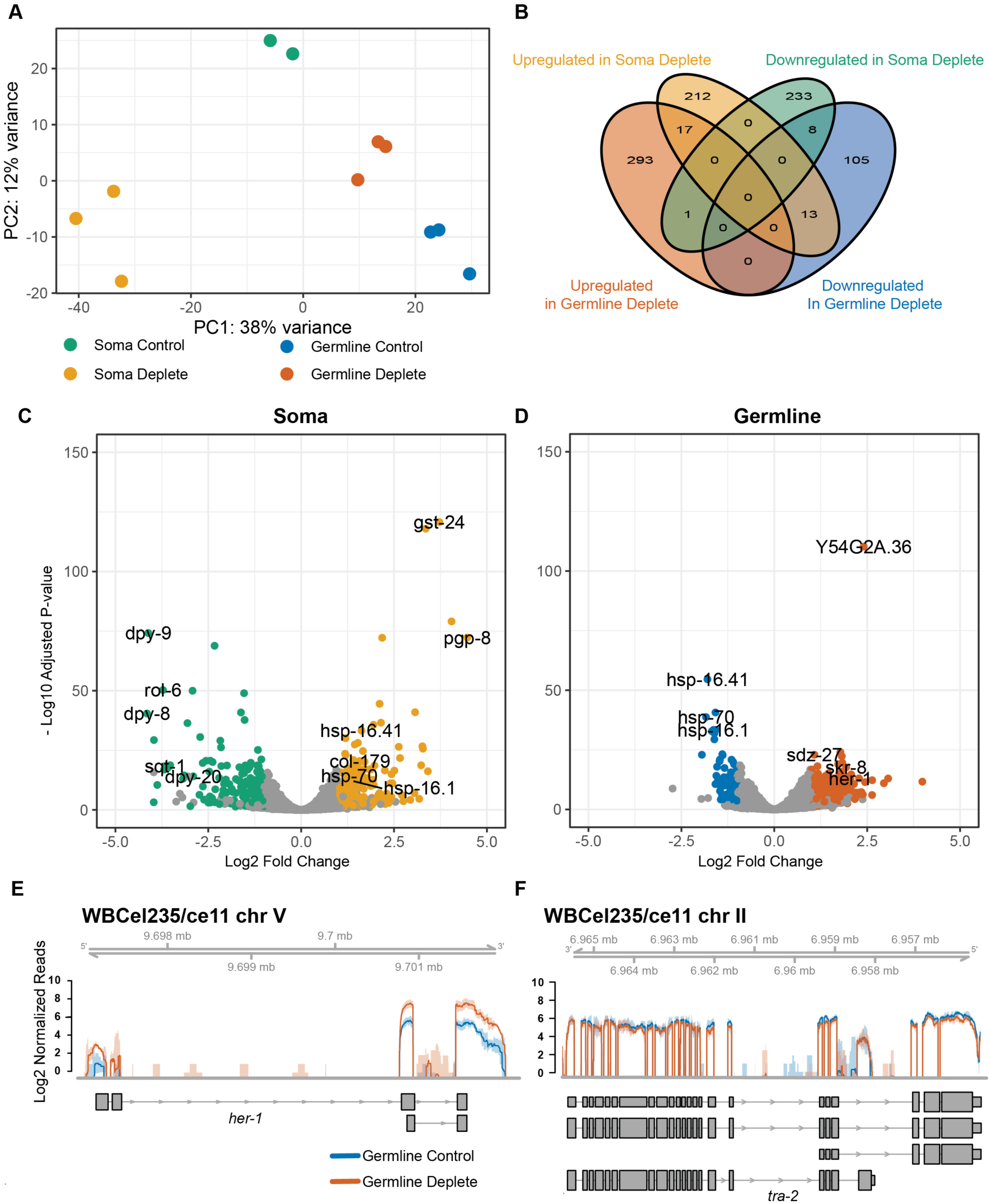
Transcriptome changes upon SACY-1 depletion. (A) PCA comparison of RNA-seq data of controls strains and the experimental samples in which SACY-1 was depleted in the germline or soma, as indicated. Three biological replicates were analyzed for each sample; however, one of the control samples for the soma depletion exhibited evidence of RNA degradation and was excluded from the analysis. (B) A Venn diagram showing the limited overlap of upregulated genes (2-fold; adjusted p<0.05, FPKM deplete > 2.5 and mean counts > 25) and downregulated (2-fold; adjusted p<0.05, FPKM control > 2.5 and mean counts > 25) genes in the RNA-seq datasets. (C, D) Volcano plots showing the log2 fold change in expression versus the −log10 of the adjusted p value of genes following SACY-1 depletion in the soma (C) or germline (D). (E, F) The normalized coverage of sequencing reads across *her-1* (E) and *tra-2* (F) following depletion of SACY-1 in the germline. The solid lines represent the mean of the biological replicates and shaded regions represent the corresponding confidence interval. Note, the pattern of *tra-2* splicing is not affected.

#### Changes in transcript abundance following SACY-1 depletion

We observed two classes of transcriptome alterations upon depletion of SACY-1 in the germline or soma: changes in transcript abundance and alterations in splicing patterns. In terms of transcript abundance, we observed 242 down-regulated genes (two-fold down-regulation, adjusted p<0.05, FPKM in soma control ≥ 2.5, mean counts across samples > 25) in the RNA samples depleted for somatic SACY-1 (Figure 6, B and C; File S2). Notably these down-regulated genes included many cuticle collagen genes and genes affecting cuticular morphology and body size (*col-17*, *col-41*, *col-46*, *col-47*, *col-90*, *col-128*, *col-149*, *dpy-3*, *dpy-4*, *dpy-5*, *dpy-6*, *dpy-8*, *dpy-9*, *dpy-13*, *dpy-20*, *lon-3*, *mlt-7*, *qua-1*, *rol-6*, *rol-8*, *sqt-1*, and *sqt-2*). Consistent with this observation, the top enriched gene ontology (GO) term for transcripts reduced in abundance in the SACY-1 soma-depleted samples was “cuticle development involved in collagen and cuticulin-based cuticle molting cycle” (Figure S7B). We also observed 242 up-regulated genes (two-fold up-regulation, adjusted p<0.05, FPKM in SACY-1 soma-deplete ≥ 2.5, mean counts across samples > 25) in the SACY-1 soma-depleted samples (Figure 6, B and C; File S2). The top enriched GO term for transcripts with increased abundance in the SACY-1 soma-depleted samples related to cellular responses to heat stress, the unfolded protein response, and innate immune responses (Figure S7), suggesting that the organism might perceive the reduction of *sacy-1(+)* function as a stressor and might then mount a response that then alters the transcriptome.

In the SACY-1 germline-depleted samples, we observed 126 down-regulated genes (two-fold down-regulation, adjusted p<0.05, FPKM in germline control ≥ 2.5, mean counts across samples > 25; Figure 6, B and D; File S2). The top enriched GO terms for transcripts with decreased abundance in the SACY-1 germline-depleted samples included the response to heat stress and the unfolded protein response (Figure S7B), suggesting the response to *sacy-1(+)* depletion differs between the soma and germline. Among the 311 transcripts increased in abundance in the SACY-1 germline-depleted sample was *her-1* (two-fold up-regulation, adjusted p<0.05, FPKM in SACY-1 germline-deplete ≥ 2.5, mean counts across samples > 25; Figure 6, B, D, and E; File S2). This might be due to an increase in X chromosome non-disjunction in embryos located in the uterus following germline depletion of *sacy-1(+)*, but this possibility was not investigated. Because *her-1* likely encodes an inhibitory ligand for the TRA-2 receptor in the sex-determination pathway (Perry *et al*. 1993; Figure 3), we tested whether the *her-1(hv1y101)* null mutation could suppress the Mog phenotype of the recessive gain-of-function *sacy-1(tn1481)* mutation, but this proved not to be the case (n=53 gonad arms). Since the genetic epistasis results suggest that SACY-1 promotes the expression of TRA-2 (Figure 3), we examined the effect of SACY-1 depletion in the germline on the levels of *tra-2* mRNA and the fidelity of its splicing. We observed no statistically significant change in *tra-2* mRNA levels or its splicing patterns (Figure 6F). We did not examine the expression of TRA-2 protein after SACY-1 depletion because a recent study suggested that TRA-2 protein expression in the wild-type germline is below the immunofluorescence detection limit of immunofluorescence (Hu *et al*. 2019).

#### Alteration of splicing patterns following SACY-1 depletion

In the SACY-1 soma-depleted samples, we observed significant (FDR<0.05) alterations in splicing patterns for 1606 transcripts (Figure 7A). These splicing alterations fell into several broad classes: the use of alternative 5’ splice sites, the use of alternative 3’splice sites, abnormal splicing within an exon, skipped exons, and retained introns. The largest class of splicing changes was the use of alternative 3’ splice sites, consistent with the fact that DDX41 was shown to be recruited to the C complex, which mediates the second step in splicing (Bessonov *et al*. 2008). Multiple splicing defects were sometimes observed within a single gene. For example, in the case of the RNA-binding protein ETR-1, which has multiple isoforms and is expressed in the soma and germline (Boateng *et al*. 2017), depletion of SACY-1 results in intron retention and multiple alterations in 3’-splice-site usage (Figure 7B). In the SACY-1 germline-depleted samples we observed significant (FDR<0.05) alterations in splicing for 796 transcripts (Figure 7A). Thus, splicing defects appeared less prevalent in the SACY-1 germline-depleted samples than the SACY-1 soma-depleted samples. One possibility is that nonsense mediated decay or other surveillance pathways actively clear misspliced mRNAs from the germline. Some alternative splicing events were observed in both the RNA preparations depleted for SACY-1 in the germline and the soma (Figure 7A). For example, we observed retention of an intron in *prdx-6* mRNA in both experiments (Figure 7C). Likewise, we observed alternative 3’ splice site selection for the heterochronic gene *lin-28* in both SACY-1-depleted samples (Figure 7D).

**Figure 7.**
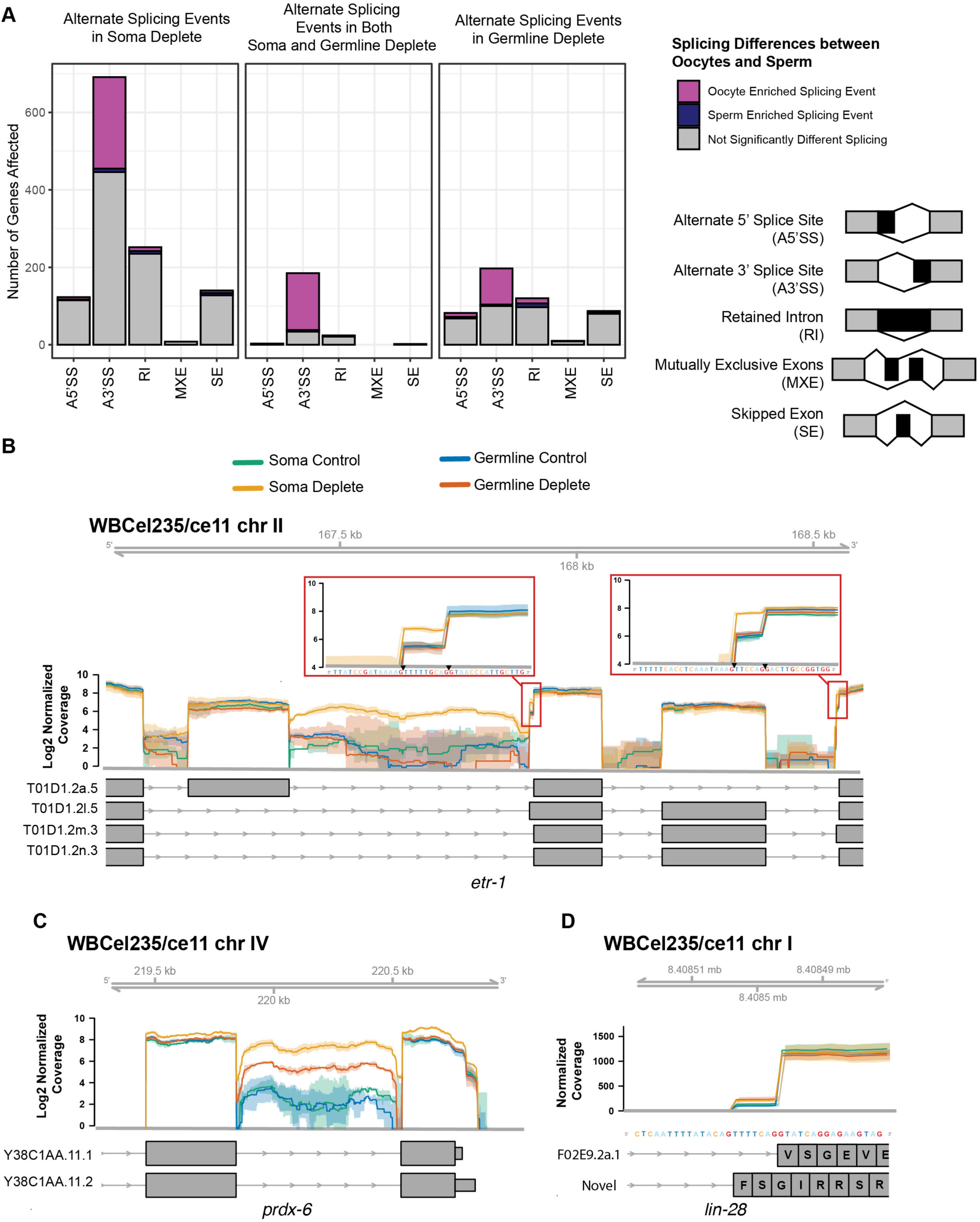
Quantification of altered splicing patterns upon SACY-1 depletion. (A) Bar graphs showing the number of genes with statistically significant (FDR<0.05) changes in splicing patterns. The legend at the right depicts the nature of the observed splicing changes: A5’SS= Alternate 5’ Splice Site, A3’SS = Alternate 3’ Splice Site, RI= Retained Intron, MXE=Mutually Exclusive Exons, SE=Skipped Exon (B–D) Examples of alterations in splicing patterns following SACY-1 depletion in the germline or soma as indicated. The *etr-1* gene shows pronounced intron retention and two alternatively spliced 3’ sites in the SACY-1 soma depleted (gold) sample (B). A subset of *etr-1* transcript annotations are shown. The *prdx-6* gene has a retained intron in the SACY-1 soma depleted (gold) and germline depleted (red) samples (C). The soma and germline depleted samples have an increase in the usage of alternate splice acceptor in the *lin-28* gene that results in an altered reading frame (D). The solid lines represent the mean of the biological replicates and shaded regions represent the corresponding confidence interval.

#### Germline sex-specific splicing patterns and the involvement of SACY-1

As we analyzed the changes in splicing patterns following SACY-1, we noticed changes in alternative splicing patterns that appeared to correlate with the sex of the germline. To investigate this observation more fully, we reanalyzed the data of Ortiz *et al*. (2014), who reported the transcriptomes of genotypically XX animals exclusively undergoing oogenesis or spermatogenesis as a consequence of containing a loss-of-function *fog-2* mutation or a gain-of-function *fem-3* mutation, respectively. This dataset was previously used to identify alternative isoform usage in the germline (Oritz *et al*. 2014). In analyzing their dataset for alternative splicing events, we observed 1600 genes for which there was a significant (FDR<0.05) germline sex-specific splicing pattern (Figure S8; File S3). We noted that upon SACY-1 depletion in either the soma or germline, oocyte-enriched splicing events were favored (Figure 7A and Figure S8). This result suggests that SACY-1 plays a role in the selection of 3’ splice sites for many genes and raises the possibility that the appropriate balance of “spliceoforms” may be play a role in cellular differentiation.

## DISCUSSION

In this study, we report the results of a molecular genetic and biochemical analysis of SACY-1/DDX41 in *C. elegans*, conducted to gain potential insights into how *DDX41* mutations might contribute to MDS and AML in humans. DDX41 is a component of the spliceosomal C complex (Jurica *et al*. 2002; Bessonov *et al*. 2008), which carries out the second step in splicing. Pre-mRNA splicing is an essential process in eukaryotes (reviewed by Wahl *et al*. 2009). Thus, the finding that mutations in highly conserved genes encoding spliceosomal components are frequently found in hematological malignancies was unexpected (reviewed by Yoshida and Ogawa 2014). Unfortunately, the genetic properties of disease-causing mutations in spliceosomal proteins have been difficult to assess. Nonetheless, the observation that many of these missense mutations map to highly conserved amino acids, within biochemically defined structural and functional domains, might suggest that they reduce but not eliminate gene function. An alternative possibility is that some of the oncogenic missense mutations might confer antagonistic gain-of-function properties. Indeed, oncogenic mutations in several splicing factors (e.g., SF3B1, U2AF1, SRSDF2, and SRSR2) present in the heterozygous condition (Yoshida and Ogaawa 2014); however, it is probable that newly arising mutations in hematopoetic lineages, in combination with the germline mutations, contribute to neoplastic development.

In this regard, the *DDX41* mutations inherited and arising in familial cases of AML are particularly informative (Polprasert *et al*. 2015). Affected individuals often inherit one copy containing a mutation, which is likely to be a null mutation (e.g., a frameshift mutation, pD140fs). In addition to this *DDX41* germline mutation, the neoplasms in these patients invariably contain a second copy with a somatic mutation that arose in the hematopoietic lineage. Several specific *DDX41* somatic mutations (e.g., R525H) arise independently with high prevalence, suggesting that they markedly contribute to neoplastic development. These somatic *DDX41* mutations are invariably missense mutations affecting conserved amino acids but have never been found to include candidate null alleles (e.g., frameshift mutations or premature termination codons). Thus, these somatically arising mutations are unlikely to eliminate DDX41 function. The presumption is that biallelic mutations disrupting DDX41 function might be lethal. In line with this view, a recent report found that two related patients with biallelic DDX41 missense mutations exhibited a more severe syndrome characterized by dysmorphic skeletal and facial features, psychomotor delays, intellectual disability, and early onset leukemia (Diness *et al*. 2018).

Here we build upon our prior work (Kim *et al*. 2012) to conduct a comprehensive molecular genetic analysis of SACY-1/DDX41 function in *C. elegans*. Recent work of others has used the *C. elegans* system to gain information on oncogenic mutations in the SF3B1 spliceosomal protein (Serrat *et al*. 2019). Our results reveal that *sacy-1* mutations confer a range of phenotypes, from highly pleiotropic defects affecting the germline and soma to very specific defects affecting cell differentiation and cell cycle regulation. Reduction-of-function *sacy-1* alleles are homozygous viable and fertile, yet affect germline sex determination and the regulation of oocyte meiotic maturation (Kim *et al*. 2012; this work). Animals homozygous for *sacy-1* null mutations grow to adulthood but exhibit a gamete degeneration phenotype and are sterile. Feminization of the germline delays oocyte degeneration, which enables *sacy-1* null mutant females to be mated and produce embryos. However, these embryos invariably die, revealing an essential maternal requirement for development. Maternally provided *sacy-1(+)* must in turn be sufficient for homozygous *sacy-1* null mutant animals (produced from heterozygous parents) to grow to adulthood.

Several *sacy-1* mutant alleles exhibit genetic properties that suggest they can counteract *sacy-1(+)* function, potentially by compromising the function of the spliceosome. Most notable among these mutations is *sacy-1(tn1481)*, which confers a masculinization of the germline phenotype resulting from the overproduction of sperm to the exclusion of the oocyte fate. Since multiple reduction-of-function *sacy-1* alleles suppress the feminization of the germline phenotype caused by null mutations in *fog-2* (Kim *et al*. 2012; this work), *sacy-1(+)* must possess a function to promote the oocyte fate. This oocyte-promoting function of *sacy-1(+)* might not be essential because *sacy-1* null mutants produce oocytes, which nonetheless undergo necrotic degeneration. The caveat here is that *sacy-1* null mutants only develop to adulthood because of maternally provided *sacy-1(+)*, which might be sufficient to promote oogenesis. In any case, the *sacy-1(tn1481)* mutation might disrupt an oocyte-promoting function either by interfering with maternally provided SACY-1 activity or proteins with which it associates.

Several observations are consistent with the possibility that *sacy-1(tn1481)* interferes with or compromises the function of the spliceosome in germline sex determination in a dosage-sensitive manner. This view is also supported by our tandem affinity purification results showing that SACY-1 is a component of the *C. elegans* spliceosome and genetically interacts with spliceosomal components. Significantly, multiple *C. elegans* spliceosomal components, functioning at different steps of the splicing reaction, can mutate to a masculinization of germline phenotype (reviewed by Zanetti and Puoti 2013). Interestingly, when placed in trans to a *sacy-1* null mutation (e.g., *tm5503*), the resulting *sacy-1(tn1481)/sacy-1(tm5503)* heterozygous animals are viable and fertile and can be maintained indefinitely as a heterozygous strain. This highly informative genetic result suggests that a single dose of *sacy-1(tn1481)* can mediate all its essential functions, but that two doses of the mutant protein interferes with the normal mechanisms of germline sex determination (i.e., *sacy-1(tn1481)* is a recessive gain-of-function antimorphic mutation). Although *sacy-1(tn1481)/sacy-1(tm5503)* heterozygous adult hermaphrodites are fertile, when *sacy-1(tn1481); fem-3(e1996)* females are mated to wild-type males, they produce embryos that invariably fail to hatch. Similar results are obtained with the six other *mog* genes (Graham and Kimble 1993; Graham *et al*. 1993), which encode spliceosomal proteins (Puoti and Kimble 1999; Puoti and Kimble 2000; Belfiore *et al*. 2004; Kasturi *et al*. 2010; Zanetti *et al*. 2011). Interestingly the P222L amino acid substitution found in *sacy-1(tn1481)* is adjacent to the Q motif, which participates in nucleotide binding and hydrolysis (Schütz *et al*. 2010). It is tempting to speculate that SACY-1/DDX41 might promote remodeling of spliceosomes during the splicing reaction and that two doses of SACY-1 P222L might disrupt these rearrangements. In a similar vein, *sacy-1(tn1479*[G504E]) exhibits a phenotype more severe than a null allele, suggesting that this mutation in the helicase domain too might possess a dosage-sensitive antimorphic activity. Whether missense alleles in human DDX41 also possess antimorphic activity will require additional work, including biochemical analyses.

We imported the oncogenic *DDX41*[R525H] mutation into *C. elegans* using CRISPR-Cas9 genome editing (R534H in SACY-1). Our analysis revealed that in *C. elegans*, this mutation possesses very weak antagonistic activity. While it is possible that the genetic properties of this mutation might differ between *C. elegans* and mammalian systems, it is worthwhile noting that the oncogenic variant must support the high levels of proliferation characteristic of neoplastic cells. While all the *sacy-1* mutations we isolated in forward genetic screens occur at conserved amino acids, none of them match the oncogenic mutations thus far isolated (Polprasert *et al*. 2015; Cardoso *et al*. 2016; Lewinsohn *et al*. 2016; Li *et al*. 2016; Diness *et al*. 2018; Sébert *et al*. 2019). This observation is consistent with the idea that the oncogenic mutations are at most weakly antagonizing or weak reduction-of-function mutations, and thus would not have been isolated in forward genetic screens that require a substantial reduction in function.

An attractive idea is that oncogenic mutations affecting the spliceosome contribute to neoplastic development through effects on gene expression occurring through alterations in RNA splicing, as well as effects on the transcriptional machinery or RNA stability (Yoshimi *et al*. 2019). Consistent with this idea, RNAi depletion of the *C. elegans* spliceosomal protein RSR-2 affects transcript levels without major impacts on splicing (Fontrodona *et al*. 2013). In this study, we examined the effects of SACY-1 depletion at the adult stage on the transcriptome. In the soma and the germline, we observed splicing-independent impacts on the abundance of many transcripts. The gene expression changes observed suggest that depletion of SACY-1 might elicit a stress response. If analogous processes occur after perturbations of the spliceosome in humans, such stress responses might contribute to oncogenesis and could represent therapeutic targets. In *C. elegans*, we observed instances of missplicing following SACY-1 depletion, though missplicing events more prevalent when SACY-1 was depleted from the soma as compared to the germline. The extent to which gene expression changes and splicing alterations contribute to the various *sacy-1* mutant phenotypes will require further study but will likely provide insights relevant to spliceosomal perturbations in humans.

## ACKNOWLEDGMENTS

This paper is dedicated to the memory of Michael A. Miller, our colleague and friend whose scientific contributions will not be forgotten. We thank Donna Coetzee for technical assistance. We are grateful to Joshua Arribere, Daniel Dickinson, Barth Grant, Tim Schedl, and Jordan Ward for providing strains or reagents. Some strains were provided by the Caenorhabditis Genetics Center, which is funded by grant P40OD010440 from the NIH Office of Research Infrastructure Programs. We thank Gabriela Huelgas-Morales, Zohar Sachs, and Todd Starich for their helpful suggestions for the manuscript. This work was supported by NIH grant GM57173 to D.G.

